# A reference library for the identification of Canadian invertebrates: 1.5 million DNA barcodes, voucher specimens, and genomic samples

**DOI:** 10.1101/701805

**Authors:** Jeremy R. deWaard, Sujeevan Ratnasingham, Evgeny V. Zakharov, Alex V. Borisenko, Dirk Steinke, Angela C. Telfer, Kate H.J. Perez, Jayme E. Sones, Monica R. Young, Valerie Levesque-Beaudin, Crystal N. Sobel, Arusyak Abrahamyan, Kyrylo Bessonov, Gergin Blagoev, Stephanie L. deWaard, Chris Ho, Natalia V. Ivanova, Kara K. S. Layton, Liuqiong Lu, Ramya Manjunath, Jaclyn T.A. McKeown, Megan A. Milton, Renee Miskie, Norm Monkhouse, Suresh Naik, Nadya Nikolova, Mikko Pentinsaari, Sean W.J. Prosser, Adriana E. Radulovici, Claudia Steinke, Connor P. Warne, Paul D.N. Hebert

**Affiliations:** Centre for Biodiversity Genomics, University of Guelph, Guelph, Ontario, Canada

## Abstract

The reliable taxonomic identification of organisms through DNA sequence data requires a well parameterized library of curated reference sequences. However, it is estimated that just 15% of described animal species are represented in public sequence repositories. To begin to address this deficiency, we provide DNA barcodes for 1,500,003 animal specimens collected from 23 terrestrial and aquatic ecozones at sites across Canada, a nation that comprises 7% of the planet’s land surface. In total, 14 phyla, 43 classes, 163 orders, 1123 families, 6186 genera, and 64,264 Barcode Index Numbers (BINs; a proxy for species) are represented. Species-level taxonomy was available for 38% of the specimens, but higher proportions were assigned to a genus (69.5%) and a family (99.9%). Voucher specimens and DNA extracts are archived at the Centre for Biodiversity Genomics where they are available for further research. The corresponding sequence and taxonomic data can be accessed through the Barcode of Life Data System, GenBank, the Global Biodiversity Information Facility, and the Global Genome Biodiversity Network Data Portal.

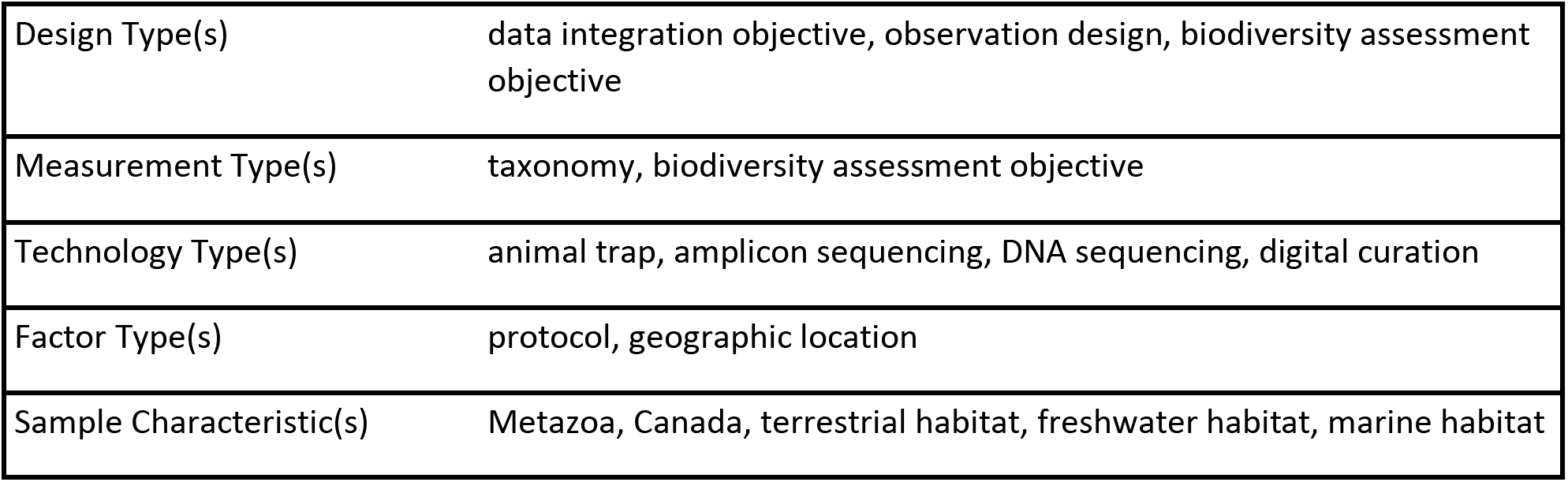

## Background & Summary

High-throughput sequencing platforms have enabled a novel approach to biodiversity surveys by making it possible to identify the array of species present in bulk collections or environmental DNA [1–3]. This ‘metabarcoding’ approach is enabling rapid assessment of the species composition of complex substrates, communities, and environments, from gut contents and feces to soil and aquatic systems, to ancient environments captured in sediments or permafrost [1,2,4,5]. The process begins by obtaining large numbers of sequence records from one or more target amplicons, or DNA barcodes [6] from the constituent organisms. This is followed by querying these sequences against a reference library derived from carefully identified records. Two reference databases are commonly employed for macro-organisms: NCBI’s GenBank [7] and the Barcode of Life Data System (BOLD; [8]). Custom reference libraries are more restrictive in taxonomic scope, but they are less prone to introducing errors in identification (e.g. [9,10]).

An adequately parameterized reference library is critical for robust taxonomic assignments (e.g. [11,12]). These libraries gain utility and reliability when each species is represented by geographically distinct populations [13,14] to capture intraspecific variation. As well, particularly for speciose genera, the libraries should ideally include multiple species [15]. The use of authoritatively identified records can provide reliable species-level assignments, but can also reveal limitations of the barcode marker(s) in particular taxa [11,13,16,17]. A significant proportion of the barcode records in reference databases are only identified to a family level [17,18] in part because some 90% of multicellular species are undescribed. Despite the lack of species-level identifications, such records are useful for assigning taxa to higher taxonomic categories [8,12,14]. While libraries with comprehensive species coverage are the ultimate goal (e.g. [19]), it is estimated that only 15% of described animal species are currently represented in public databases [20]. There is, however, a strong prospect that coverage will rise rapidly with the introduction of high throughput sequencing (HTS) protocols which come with lower analytical costs [21–23], and can recover sequences from museum and type specimens [24].

While the reliability of identifications generated through metabarcoding studies depends mainly upon access to a well-parameterized reference library, the quality of the libraries is reinforced by access to the voucher specimens, to high-quality images of these specimens, and to genomic DNA samples from them. The importance of retaining voucher specimens in systematic and ecological studies is well-appreciated [25–28] – they allow future taxonomic examination which in turn, can prevent the proliferation of incorrect identifications or ‘error cascades’ [29]. Because the retention of specimens used as a source for DNA has not been required for sequences submitted to GenBank, it contains many sequences derived from specimens which were discarded, destroyed, or misidentified [27]. Moreover, corrections to sequences assigned to the wrong species in GenBank can only be made by the data submitter, a serious impediment to curation [30]. Curation of reference libraries is also facilitated by high-resolution images of specimens, either as ‘e-vouchers’ [31] that can be digitally distributed, or better yet, images are taken in addition to voucher specimens. It is also apparent that cryopreserving genomic DNA extracts derived from the construction of reference libraries will become increasingly important [32,33]. DNA vouchering provides another opportunity for taxonomic ‘quality control’, in those taxa where the primary barcode region fails to differentiate closely-related taxa, as additional sequence information can be gathered (e.g., 16S rRNA; [5]). In fact, the retention of DNA extracts provides a basis for ‘upgrading’ records if the barcode standard eventually adopts new approaches in groups, such as plants, where data standards have changed [34], or for sequencing the entire genome [35].

This data resource presents a curated DNA barcode reference library for a substantial fraction of the Canadian invertebrate fauna, illustrates the workflows involved in its construction, and describes the resources resulting from this effort. The library was constructed and curated for over a decade, beginning with collecting efforts at sites across Canada, followed by specimen processing and DNA barcoding, and finally, taxonomic identification and validation. The library includes barcode records for 1.5 million specimens representing nearly 65,000 species, all supported by voucher specimens, digital images, and DNA extracts available for follow-up studies. Despite a national focus, we anticipate it will have wide utility in metabarcoding and related studies, due to the diversity of taxa included.

## Methods

The curated DNA barcode reference library presented here for Canadian invertebrates was constructed by a series of workflows that generate diverse products and accessible resources (Figure 1).

**Figure 1.**
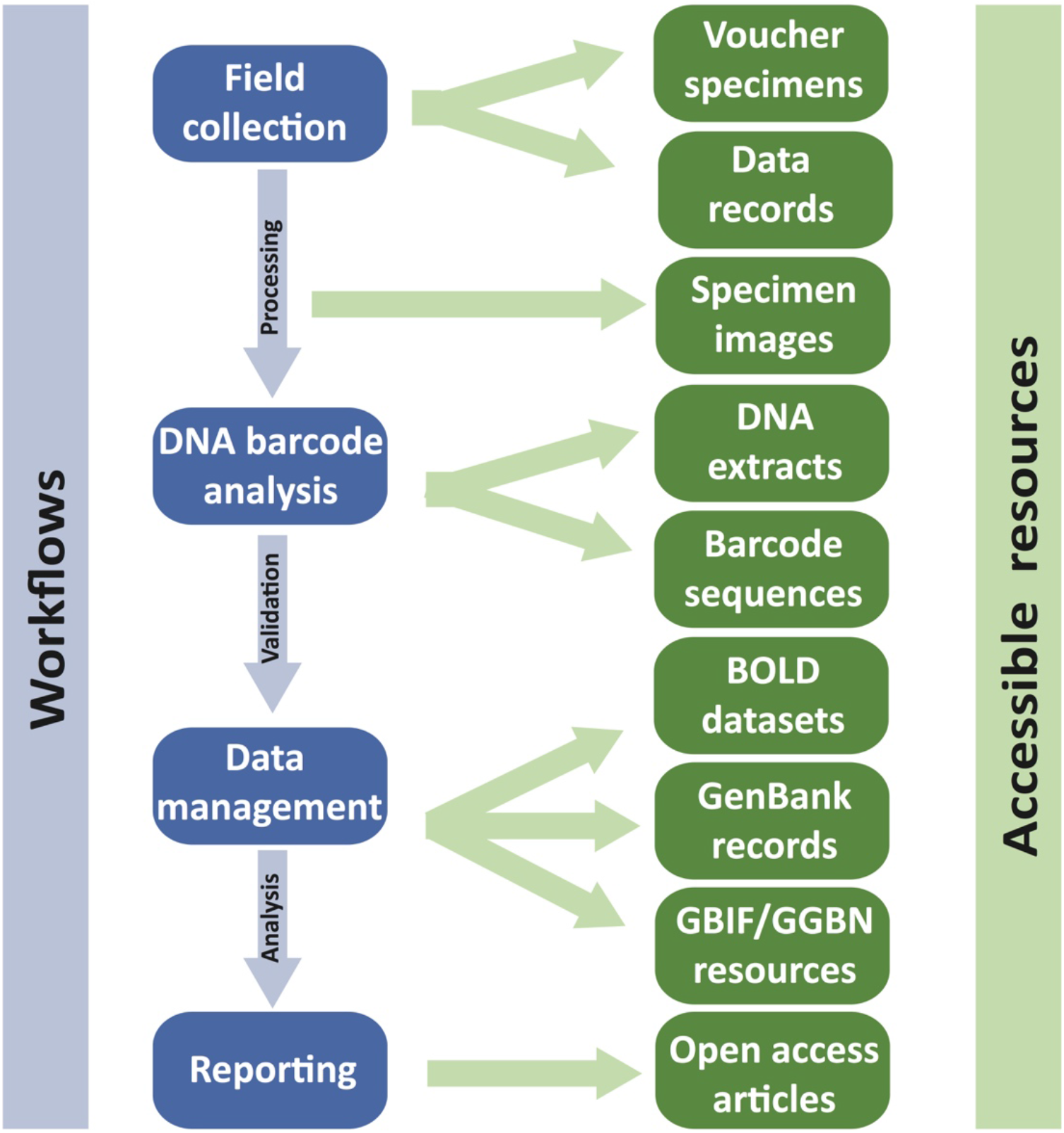
Overview of the study design to create and maintain the curated reference DNA barcode library for Canadian invertebrates.

### Field Collections

All collecting was done in compliance with national and provincial regulations and appropriate permits were obtained where necessary. No permits are required to collect invertebrates from public areas in Canada, except for species at risk; none of the collecting efforts targeted endangered or protected species. Collections of invertebrates within Canadian National Parks were made under permits NAP-2008-1636 (2008-2010), PC-2012-11074 (2012-2014), and NAP-2015-19000 (2015-2017) granted by Parks Canada. Permits and/or permissions were also obtained for invertebrate sampling in provincial parks (e.g. BC provincial parks in 2014 – British Columbia Ministry of Environment # 107242), municipal parks, conservation authorities, research properties, and other protected areas (e.g. Nature Conservancy of Canada properties).

### Sampling Localities

Specimens were collected from 3413 locations (Figure 2A), representing all 13 provinces and territories of Canada, and all 23 of its terrestrial and aquatic ecozones [36]. Collecting sites included national parks, provincial parks, municipal parks, conservation reserves, research reserves, other protected areas, school grounds, as well as industrial and residential properties. Of the 47 National Parks, National Park Reserves, and National Urban Parks in Canada, collecting was undertaken in 43 (Figure 2B). In total, 1,500,003 specimens were collected and underwent DNA barcoding (see below), of which 1,002,170 were collected within national park boundaries (herein, the ‘National Parks subset’) and 497,833 elsewhere (the ‘Other Localities subset’) (Figure 3).

**Figure 2.**
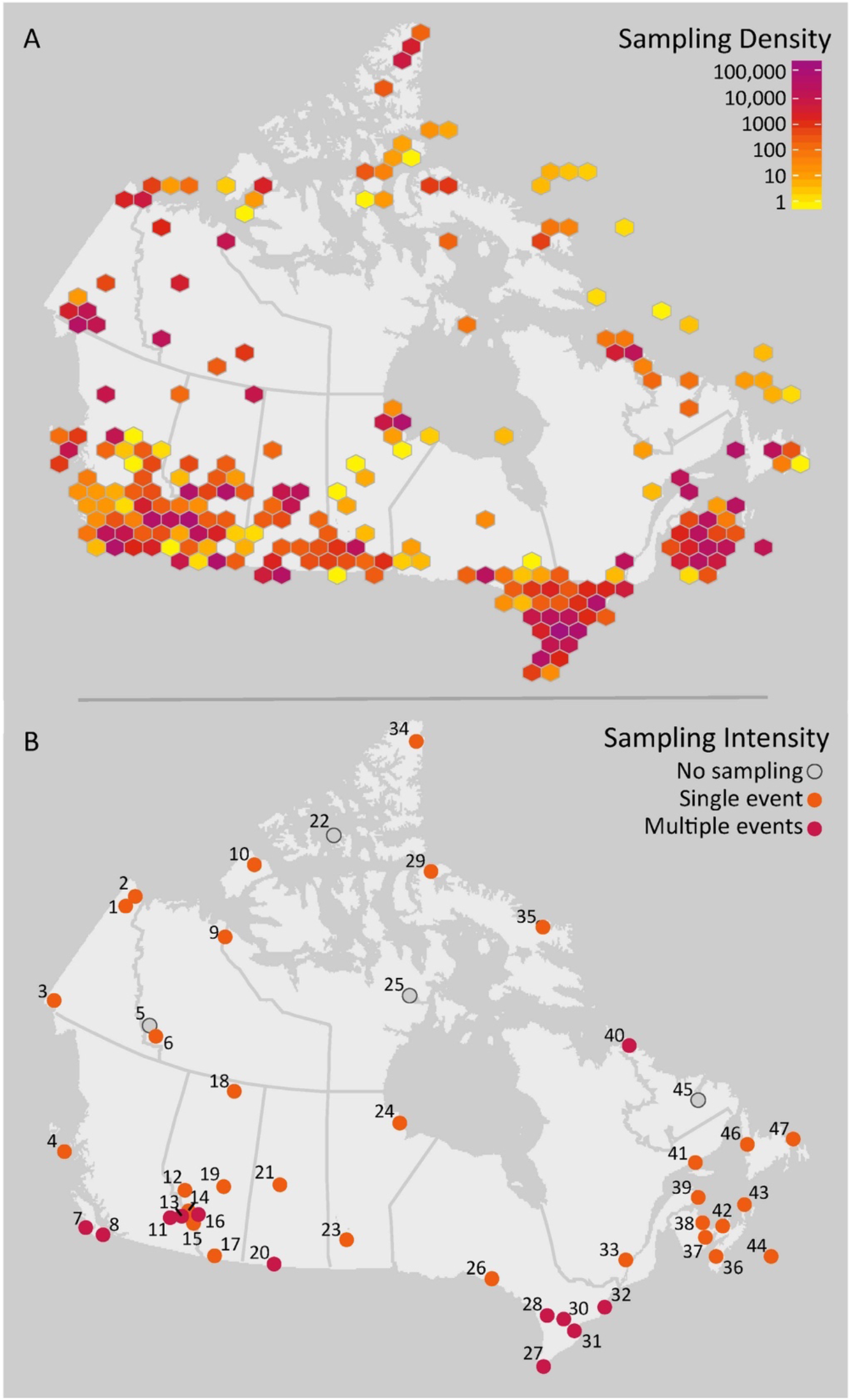
Geographic coverage of the Canadian specimen data release. A) The sampling density for the complete dataset of 1,500,031 specimen records. B) The sampling intensity for the 47 National Parks, National Park Reserves, and National Urban Parks of Canada. Numbers correspond to Parks in Table 2.

**Figure 3.**
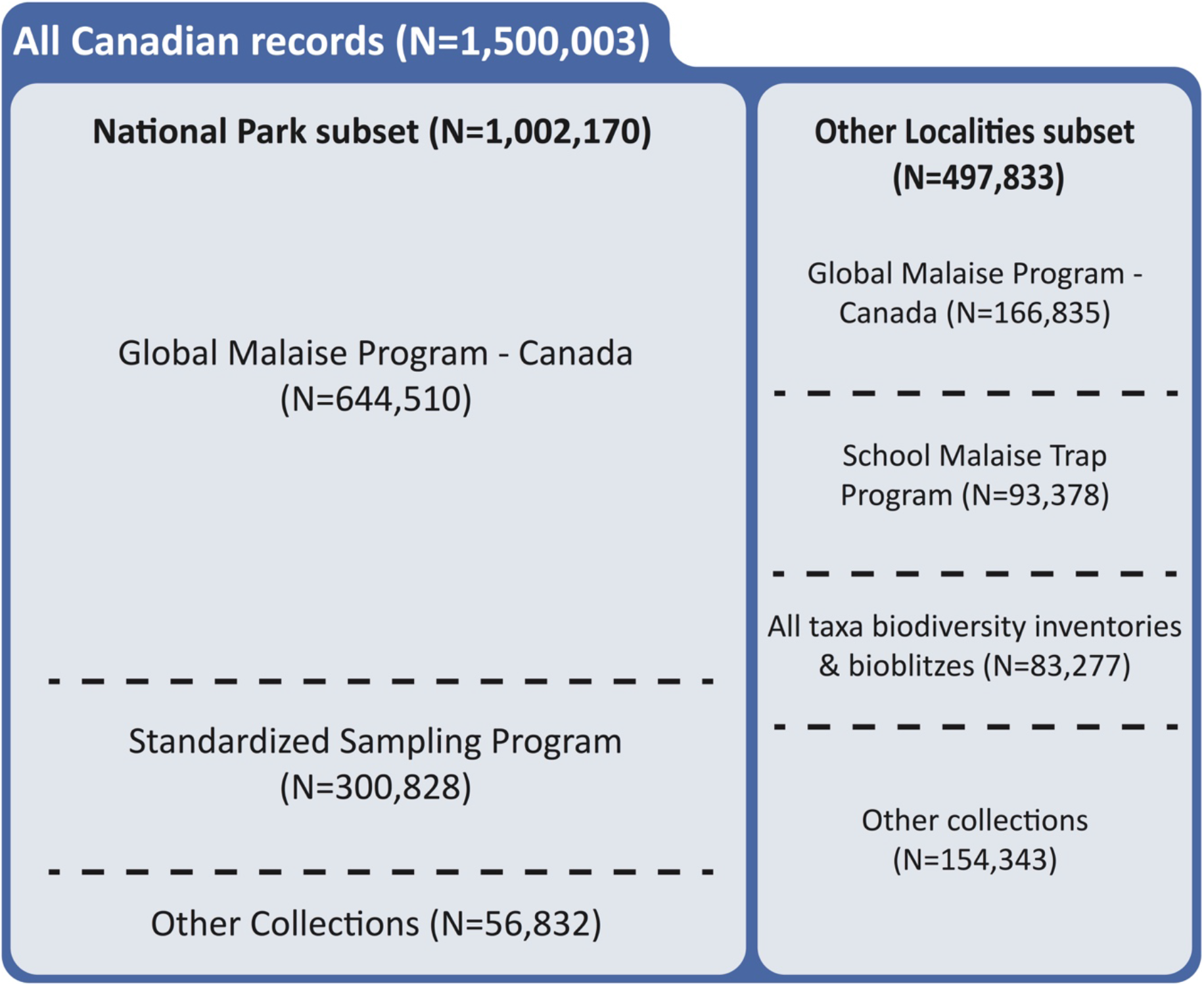
Breakdown of the Canadian specimen data release by sampling program. N = number of records.

### Sampling Methods and Programs

The 1,500,003 specimens included in this data release were obtained by both active and passive collecting methods, and as part of various sampling programs across Canada (Figure 2A, details below; Figure 2B). Specimens were predominantly collected between 2008 and 2017, but a small fraction (2.0%) was donated to the CBG from external collections dating back to 1981. The primary techniques employed for collection were Malaise traps (N=1,096,898), sweep netting (N=87,910), flight-intercept traps (N=69,162), pan traps (N=67,225), pitfall traps (N=43,771), ultraviolet (UV) light collections (N=24,501) and manual collecting (N=19,334).

#### National Parks subset

With support from Parks Canada, invertebrate surveys were conducted in 43 National Parks and National Park Reserves during 2012-2014; two thirds of the specimens (N=644,510) were collected using Townes-style Malaise traps [37–39] as part of the Global Malaise Trap Program (GMP) – Canada (www.globalmalaise.org; [40]). Most parks were only sampled during one year, but 12 were sampled in two or more years (Figure 2B). The length of the collecting season for each park spanned most of the insect flight period, but was also determined by weather conditions and, for remote locations, by accessibility of the traps for servicing (see Supplementary File 1). Malaise traps were typically serviced weekly by Centre for Biodiversity Genomics (CBG) or Parks Canada staff by replacement of the sampling bottle with a bottle containing fresh preservative.

Additional specimens in the ‘National Parks subset’ (N=300,828) were collected as part of the CBG’s Standardized Sampling Program. It ran from 2012-2014 in 23 National Parks, National Park Reserves, and National Urban Parks and employed a standard set of sampling methods that targeted a wider diversity of invertebrate fauna than Malaise traps [41]. In each Standardized Sampling (SS) locality, three representative sites were chosen based on a variety of biotic and abiotic factors, such as habitat type, vegetation, and elevation. The protocol was implemented over a one-week period and involved the deployment of a standard array of traps at each site: 1-2 Malaise traps, 1 flight-intercept trap, 10 pan traps, and 10-20 pitfall traps. In addition, 1-3 substrate samples were taken for Berlese funnel extraction, and a total of 60 min of sweep netting was performed over the week.

The remaining 56,832 (5.7%) specimens were obtained through opportunistic collecting in terrestrial, freshwater, and marine habitats using UV lights, dip nets, plankton nets, sieves, aspirators, mustard extraction, and freehand collecting.

#### Other Localities subset

The remaining 497,833 specimens in this subset were obtained through various methods and collection programs:

- the largest proportion of this subset (N=166,835) was obtained through Malaise trapping at 35 additional protected areas (not national parks) for GMP – Canada [40], including sites in proximity to ports in Vancouver, Montreal, Toronto, and Halifax (N=71,747).
- an educational program – the School Malaise Trap Program (SMTP; [42]) – deployed Malaise traps on school grounds across Canada, contributing a significant number of specimens (N= 93,378).
- all taxa biodiversity inventory (ATBI) and bioblitz [43] activities combined to provide a large number of specimens (N=83,277) using a diverse repertoire of methods. These included an ATBI at Churchill, MB from 2006 to 2009 (N=41,449) [44–47]; a long-term inventory of the *rare* Charitable Research Reserve in Cambridge, ON (N=41,608) [48]; and bioblitz [43] events involving the Ontario BioBlitz (www.ontariobioblitz.ca) and Bioblitz Canada (www.bioblitzcanada.ca) programs.
- Other collections (N=154,343) were likewise made with a variety of techniques, including our SS technique at protected areas (N=22,943)

A summary of the collection method(s) used for each program is provided in Supplementary File 2. The collection method for each specimen is included in the data resources, as well as information on trap type, weather conditions, habitat, and any deviations from the normal collection protocol when this information is available. These programs covered 39 protected areas, including provincial parks, municipal parks, conservation reserves, ecological reserves, research reserves, and Nature Conservancy of Canada properties. Adding the collections in 43 national parks, a total of 1,132,347 occurrence records were derived from 82 protected areas across Canada.

### Code Availability

The Barcode of Life Data System (BOLD; www.boldsystems.org; [8]) was used as the primary workbench for creating, storing, analyzing, and validating the specimen and sequence records and the associated data resources [48]. The BOLD platform has a private, password-protected workbench for the steps from specimen data entry to data validation (see details below), and a public data portal for the release of data in various formats. The latter is accessible through an API (http://www.boldsystems.org/index.php/resources/api?type=webservices) that can also be controlled through R [49] with the package ‘bold’ [50].

### Specimen Processing and DNA Barcode Analysis

The CBG has an efficient workflow for collecting, sorting, processing, and DNA barcoding specimens for reference library construction. As detailed protocols are outlined in other publications [51,52], only summary details are provided here (Figure 1).

Following collection, and prior to sorting, bulk samples and specimens were stored in −20°C freezers, remaining in or transferred to 95% ethanol. All specimens from a trap sample or collection event were prepared for DNA barcoding, except in those cases where initial inspection suggested the presence of a very large number of specimens of a particular species. In these cases, 5 to 95 representatives of each morphospecies were prepared for sequence analysis and excess specimens were retained in ethanol at −20°C. Larger specimens were pinned and one leg was removed for DNA extraction; smaller specimens were placed directly into 95% ethanol in either a) a sample tube rack, where a leg was later tissue-sampled for DNA extraction or b) a microplate, where the entire specimen was used for DNA extraction with an added step of recovering exoskeletal remains after non-destructive lysis and DNA extraction (‘voucher recovery’, [53]).

Subsequent barcode analysis was performed following standard methods [52,54]; the stages include tissue lysis, DNA extraction, PCR amplification of the 658 base pair (bp) fragment of the cytochrome *c* oxidase subunit I (COI) gene, cycle sequencing, and subsequent Sanger sequence analysis. The resultant sequences, as well as electropherograms, and primer details for all specimens were uploaded to BOLD.

### Barcode Index Numbers

For all sequences uploaded to BOLD, the records were assigned operational taxonomic units called Barcode Index Numbers (BINs) by the Refined Single Linkage (RESL) algorithm implemented on BOLD [55]. Individual records are either assigned to an existing BIN or found a new BIN, but they only enter the RESL analysis if they meet the following criteria: greater than 300 bp coverage of the barcode region, less than 1% ambiguous bases, and no stop codon or contamination of the sequence. For inclusion into an existing BIN, sequence records must include >300 bp of the barcode region (between positions 70 and 700 of the BOLD alignment) while records that establish a new BIN must include >500 bp of the barcode region. The RESL algorithm runs monthly on all qualifying barcode sequences in BOLD – which currently contains 6.9 million animal specimen records and 0.63 million BINs (July 2019). BIN designations and assignments generated by RESL on BOLD are accessible for independent validation through the ‘BIN pages’ that aggregate the specimen and sequence information of its members (e.g., the deer tick, *Ixodes scapularis* Say: http://dx.doi.org/10.5883/BOLD:AAA1270).

### Taxonomic Assignment

Prior to processing, most specimens were identified to an order level based on morphology. Following a record’s assignment to a BIN, if that BIN contained specimens identified to a single family, genus or species, it received this identification. In cases of taxonomic discordance, the identification was applied above the level of disagreement. For example, if a BIN containing two members had one specimen assigned to genus A and the other to genus B, but both belonged to family C, the specimen would only be identified to the family level.

For specimens without a BIN assignment or where the taxonomy associated with the BIN was only to a family level, specimen sequences were compared to the complete reference library on BOLD using its Identification (BOLD-ID) Engine (available at http://v4.boldsystems.org/index.php/IDS_OpenIdEngine). A list of the top 99 sequence matches for each specimen was returned, and the taxonomy was applied where present and without discordance (as in BIN taxonomy assignment described above). Species-level identifications were assigned at ≥ 98.5% sequence similarity, genus-level identifications at ≥ 95% similarity, and family-level identifications at ≥ 90% similarity.

Specimens still lacking an identification at the family level were placed into a Neighbor-Joining tree of identified records in the same order, constructed on BOLD (see Supplementary File 3 for an example). If an unnamed specimen fell within a distinct haplogroup cluster, the lowest taxonomic level of agreement was applied to the specimen. If this approach was also unsuccessful, specimens were identified morphologically where possible, either by in-house experts or through loans to taxonomic specialists (e.g. Canadian National Collection of Insects, Arachnids, and Nematodes; Smithsonian Institution’s National Museum of Natural History).

### Specimen, DNA, and Image Storage

All voucher specimens in the dataset were archived in a secure, microclimate-controlled Specimen Archive (BIOUG). All specimen provenance data, timing of processing, and storage locator information were digitized in a custom-designed institutional database (see Technical Validation below) to allow the efficient pre-laboratory processing, data submission, archival storage, and retrieval of specimens. All vouchers are available for loan for further research, and the data are accessible in various data portals (see Data Records below).

The DNA extracts produced during barcode analysis are stored within a DNA Archive, either in −80°C freezers or dried in a trehalose or PVA-based cryoprotectant [56] and held in −20°C freezers. Information on these DNA extracts is stored in a MS Access database. Tracking of the DNA extracts through the DNA barcoding analytical steps was also captured by a custom-built PostgreSQL-based Laboratory Information Management System (BOLD-LIMS). The data necessary for the preparation of the specimen core and GGBN extension files were exported from the DNA Archive database and BOLD (see Data Records below).

Representatives of each BIN were photographed to build a digital image library to aid taxonomic validation. Specimens were photographed at high resolution and the images were made accessible through both the specimen and BIN pages on BOLD under Creative Commons (BY-NC-CA) license.

## Data Records

### Records Summary

Although the specimens were sourced from localities spanning ~4500 km in latitude and ~7000 km in longitude, sampling coverage was strongest in southern Canada (Figure 2A). Sampling coverage varied between 13 provinces and territories more than 20-fold, with N=13,225 (0.9%) for Nunavut versus N=425,049 (28.3%) for Ontario. Most of the specimens (~98%) were from terrestrial habitats followed by freshwater (~1.5%) and marine (~0.5%) environments.

Most specimens associated with this data release are available for loan or further study in the Centre for Biodiversity Genomics Collection (BIOUG). A small percentage (2.3%) of specimens were damaged or lost during processing but, in nearly all cases, other representatives of that BIN were recovered. In total, 210,585 (14.0%) specimens were photographed and these images can be accessed on both the individual specimen and BIN pages. Most BINs (N=58,126; 90.5%) in the data release are represented by an image of at least one voucher. When paired with Neighbor-Joining (NJ) trees, these images are critical for taxonomic validation and identification refinement (see Supplementary Files 3 and 4 for a NJ tree and associated images for one group of Canadian net-winged insects). The image library may also be useful as a training dataset for machine learning algorithms designed for specimen identification utilizing images (e.g. [57]).

This data release is taxonomically extensive as it includes representatives for 14 phyla, 43 classes, 163 orders, 1123 families, and 6186 genera. A very high proportion of the specimens have taxonomic assignments at the family (99.9%) and genus (69.5%) levels, but fewer (N=571,902; 38.1%) could be assigned to a species (Table 1). Of the 1,500,003 specimens included in the resource, 1,457,334 (97.2%) were either placed into an established BIN on BOLD or founded a new one, for a total of 64,264 BINs. As a proxy for species, this BIN total represents a substantial gain for the Canadian species inventory. The last thorough compilation for all invertebrates [58,59] indicated only 41,941 Canadian species and an estimated fauna of 78,821 species. Similarly, the more recent compilation of all terrestrial invertebrates by Langor [60] assembled 44,100 described species with 27,000-42,600 remaining undiscovered and/or undescribed. Flies (Diptera) dominate both specimens (N=875,215; 58.3%) and BINs (N=27,525; 42.8%) in the current reference library, followed by bees, wasps, ants and allies (Hymenoptera) and moths and butterflies (Lepidoptera) (Table 1). The ‘Other Localities subset’ included 26.9% more families although it included half as many specimens as the Parks dataset. Taxonomic resolution (measured at the species level) also varied slightly between the subsets with the ‘National Parks subset’ at 35% identified to a species versus 44% for the ‘Other Localities subset’. This variation in resolution is apparent between taxonomic categories as well; just 5-16% of mites and ticks (Acari) have a species assignment versus 99-100% for spiders (Araneae) and 82-87% for moths and butterflies (Lepidoptera).

**Table 1.**
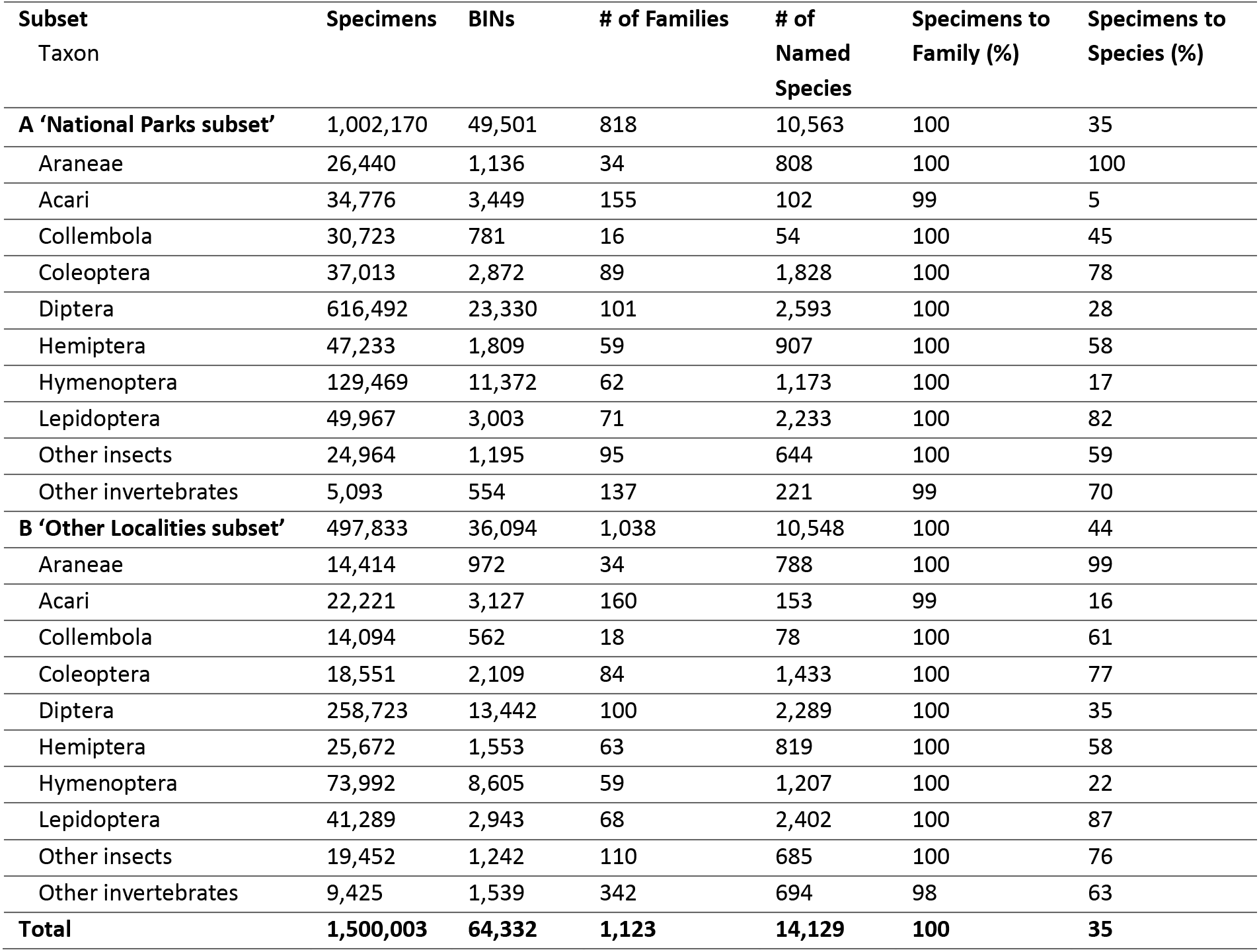
Summary data by major taxon represented within the dataset.

| Subset<br>Taxon | Specimens | BINs | # of Families | # of<br>Named<br>Species | Specimens to<br>Family (%) | Specimens to<br>Species (%) |
| --- | --- | --- | --- | --- | --- | --- |
| **A 'National Parks subset'** | 1,002,170 | 49,501 | 818 | 10,563 | 100 | 35 |
| Araneae | 26,440 | 1,136 | 34 | 808 | 100 | 100 |
| Acari | 34,776 | 3,449 | 155 | 102 | 99 | 5 |
| Collembola | 30,723 | 781 | 16 | 54 | 100 | 45 |
| Coleoptera | 37,013 | 2,872 | 89 | 1,828 | 100 | 78 |
| Diptera | 616,492 | 23,330 | 101 | 2,593 | 100 | 28 |
| Hemiptera | 47,233 | 1,809 | 59 | 907 | 100 | 58 |
| Hymenoptera | 129,469 | 11,372 | 62 | 1,173 | 100 | 17 |
| Lepidoptera | 49,967 | 3,003 | 71 | 2,233 | 100 | 82 |
| Other insects | 24,964 | 1,195 | 95 | 644 | 100 | 59 |
| Other invertebrates | 5,093 | 554 | 137 | 221 | 99 | 70 |
| **B 'Other Localities subset'** | 497,833 | 36,094 | 1,038 | 10,548 | 100 | 44 |
| Araneae | 14,414 | 972 | 34 | 788 | 100 | 99 |
| Acari | 22,221 | 3,127 | 160 | 153 | 99 | 16 |
| Collembola | 14,094 | 562 | 18 | 78 | 100 | 61 |
| Coleoptera | 18,551 | 2,109 | 84 | 1,433 | 100 | 77 |
| Diptera | 258,723 | 13,442 | 100 | 2,289 | 100 | 35 |
| Hemiptera | 25,672 | 1,553 | 63 | 819 | 100 | 58 |
| Hymenoptera | 73,992 | 8,605 | 59 | 1,207 | 100 | 22 |
| Lepidoptera | 41,289 | 2,943 | 68 | 2,402 | 100 | 87 |
| Other insects | 19,452 | 1,242 | 110 | 685 | 100 | 76 |
| Other invertebrates | 9,425 | 1,539 | 342 | 694 | 98 | 63 |
| **Total** | **1,500,003** | **64,332** | **1,123** | **14,129** | **100** | **35** |

A closer examination of the ‘National Parks’ subset reveals the recovery rate and overall complexity of the barcode-based workflow. In total, 1,148,787 specimens were processed from collecting events in these sites, but just 1,002,170 (87.2%) qualified for inclusion in the data release for four reasons. Firstly, 132,933 (11.6%) specimens were not successfully sequenced, with the order Hymenoptera comprising the largest proportion of failures (N=46,103 failed specimens; recovery rate = 73.7%), followed by Diptera (N=34,161; 94.8%), Acari (N=17,189; 66.9%), and Hemiptera (N=14,203; 76.9%). Secondly, ten sequenced records contained stop codons, indicating that a pseudogene was likely sequenced instead of the COI barcode region; their low incidence (0.001%) indicates that nuclear mitochondrial pseudogenes (NUMTs; see [61]) rarely complicate the recovery of COI through Sanger sequencing, likely because the copy number of NUMTs is far less. Thirdly, 4,799 were flagged as possible contaminations or misidentifications. Fourthly, 6,100 specimens were excluded because their sequence was either <300 bp, had >1% ambiguous bp in the barcode fragment, or they lacked both a BIN and a family assignment. And lastly, as part of the taxonomic assignment workflow, 2,737 specimens were permanently transferred to other institutions so their vouchers are unavailable at the CBG.

Because collecting efforts in the national parks varied in frequency and length (Figure 2B, Supplementary File 1), there was considerable variation in the number of BINs and specimens captured per park (Table 2, Figure 4). Values ranged from a low of 77 BINs and 715 specimens at Auyuittuq National Park to 6,806 BINs and 48,405 specimens at Jasper National Park, with an average of 2,988 BINs and 23,3017 specimens per park. By comparison, in the ’Other Localities’ subset, GMP - Canada captured 15,879 BINs and 166,835 specimens, SMTP captured 8,878 BINs and 93,378 specimens, and the combination of ATBIs and bioblitzes captured 10,721 BINs and 83,277 specimens. The sampling methods employed at each national park differed in some cases as well, further contributing to the disparity. As expected, these sampling methods each captured a differing subset of the local fauna, but in combination, they led to more comprehensive collections (Supplementary File 5).

**Table 2.**
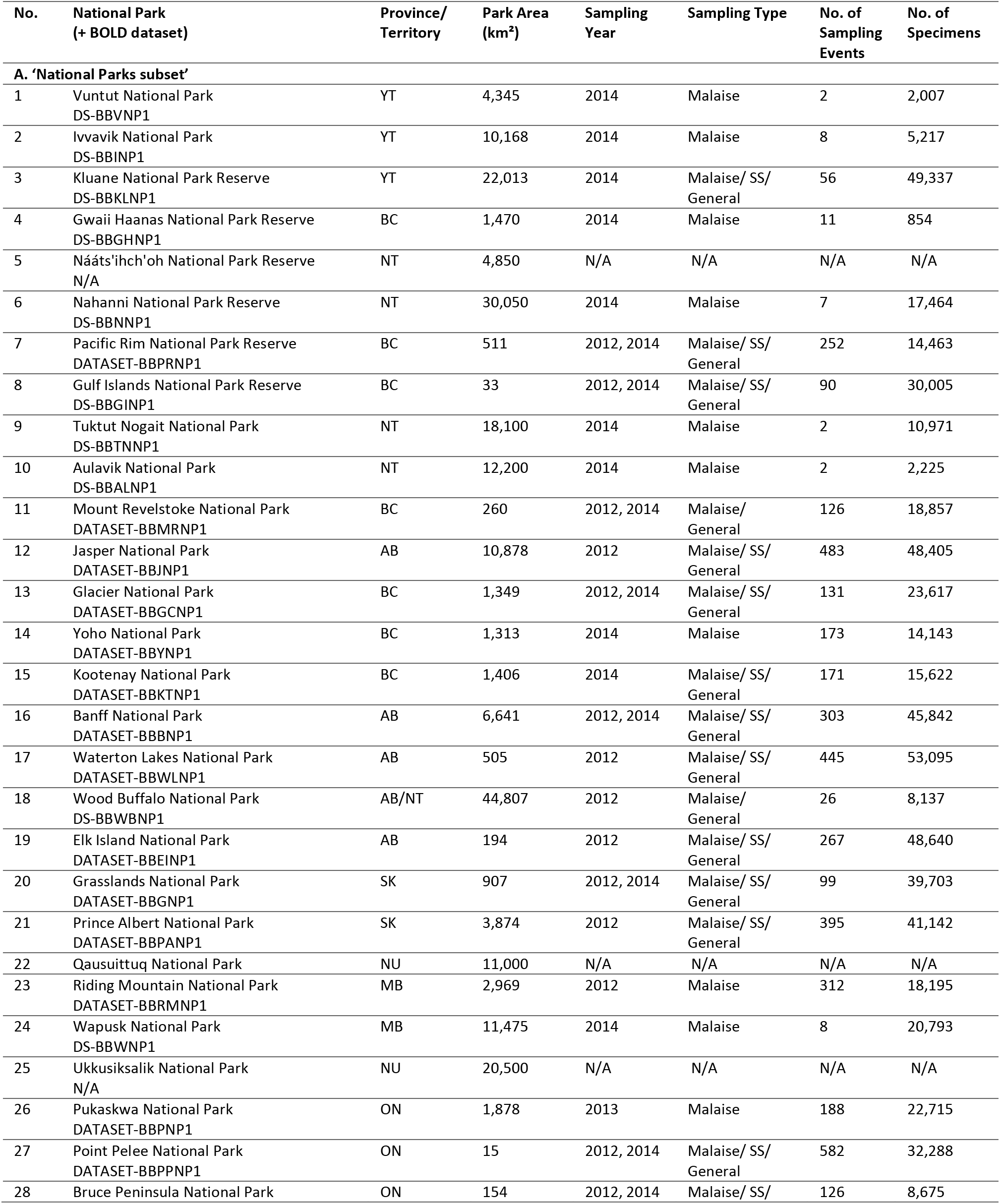
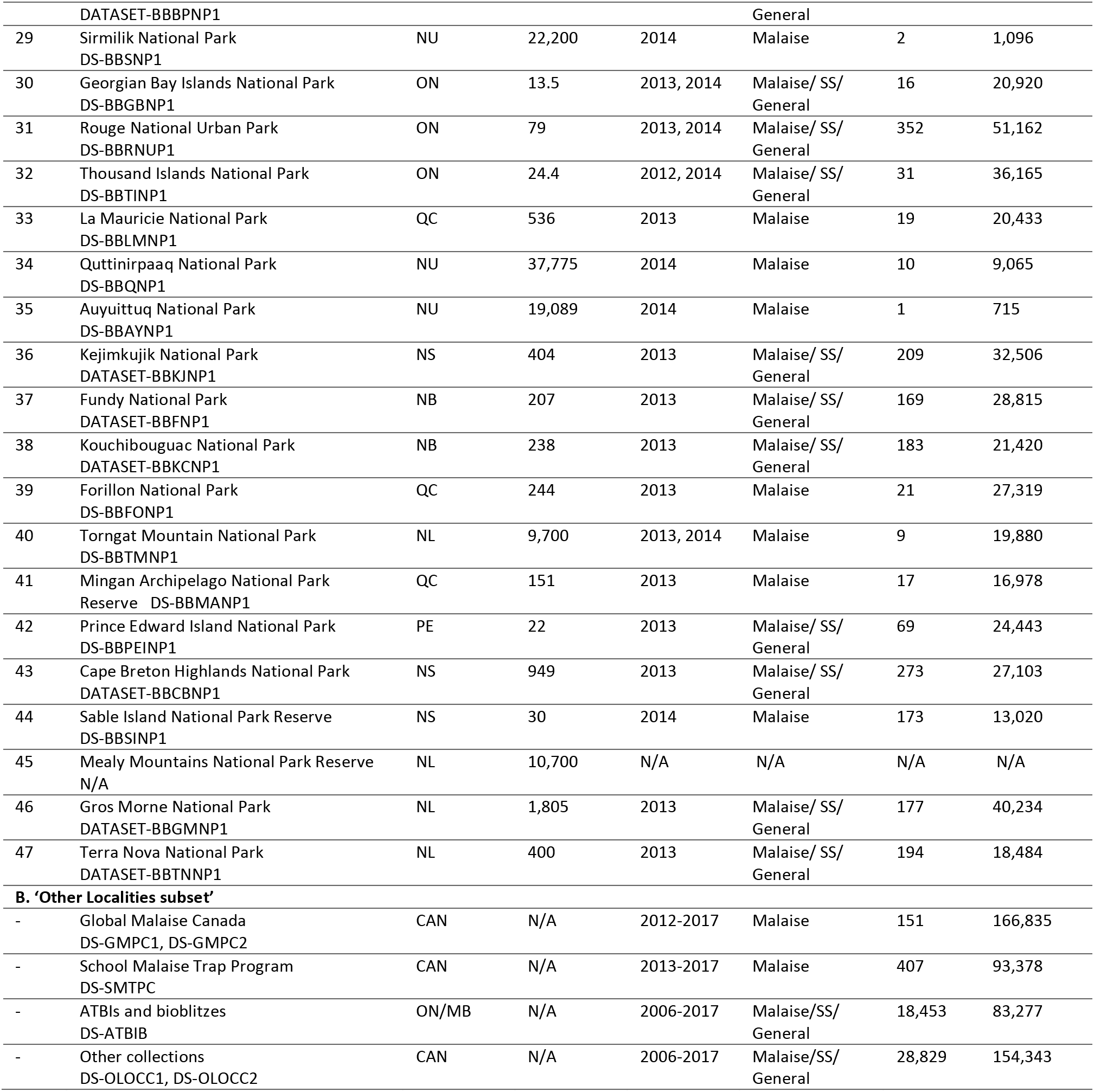
Summary data and BOLD datasets for each national park and other locality subsets. Park numbers correspond to those in Figure 2.

**Figure 4.**
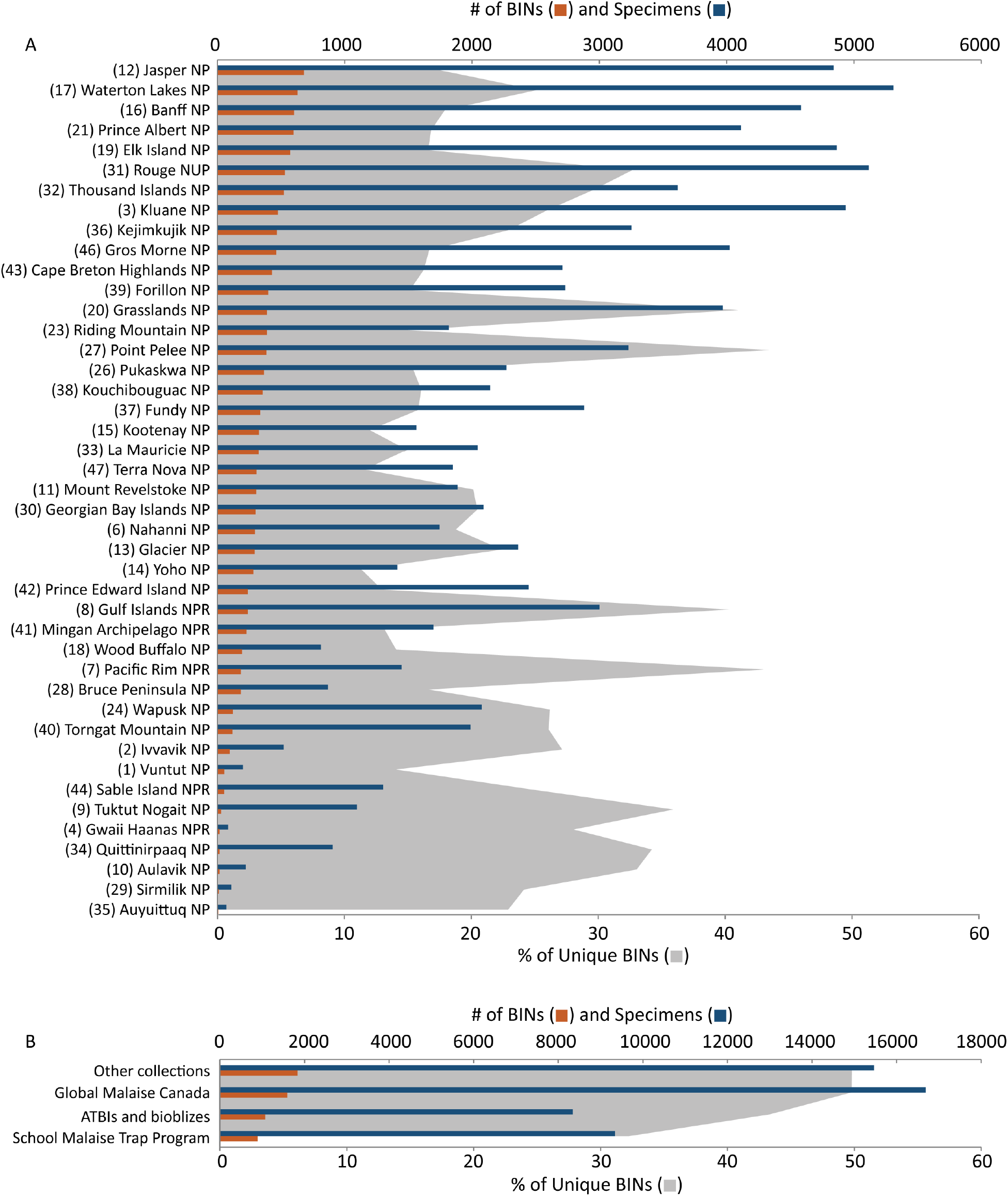
Specimen and BIN summaries for the two subsets in this data release. A) National Park subset. Numbers correspond to those in Figure 2 ranked by BIN count, and B) Other localities subset. Numbers in parentheses correspond to Parks in Figure 2. ‘Unique BINs’ refer to BINs collected only in that specific national park or collecting program.

### Records Access

The specimen and sequence data for all 1,500,003 records are available on BOLD in public datasets (see list in Table 2, where specimens are grouped by national park and major collection programs; Figure 3). The record for each specimen includes its date and locality of collection, its taxonomic assignment, and voucher specimen details. The record also includes trace files, quality scores, nucleotide sequence for the COI barcodes, and corresponding GenBank accession numbers. Condensed versions of the ‘National Parks’ and ‘Other Localities’ subsets are available in Supplementary Files 6 and 7, respectively. As noted earlier, 210,585 (14.0%) of the records possess a photograph of the specimen, nearly all with the Creative Commons Attribution-NonCommercial 4.0 (CC-BY 4.0) License. Each specimen record has been publicly released and is searchable in the Public Data Portal on BOLD (www.boldsystems.org/index.php/Public_BINSearch) or downloadable by utilizing BOLD’s API (www.boldsystems.org/index.php/resources/api). Additionally, BOLD users can log in and search for any specimen(s) from the BOLD Workbench (http://www.boldsystems.org/index.php/Login/page). BOLD’s various methods of delivering the data permit a wide range of queries and subsequent analyses (Data Citation 1).

All sequences in this data release have been submitted to GenBank. A full list of GenBank Accessions for the ‘National Parks’ and ‘Other Localities’ subsets are available in Supplementary Files 6 and 7, respectively. From the GenBank homepage (https://www.ncbi.nlm.nih.gov/genbank/), accessions can be searched as a comma-separated list. The entire dataset can be accessed through the NCBI’s BioProject PRJNA472144 (www.ncbi.nlm.nih.gov/bioproject/472144) (Data Citation 2).

After final validation, specimen data were uploaded to the Global Biodiversity Information Facility (GBIF; http://www.gbif.org) as a Darwin Core Archive [62]. The data are available from the University of Guelph’s installation of the Integrated Publishing Toolkit [63] and webserver (https://ipt.uoguelph.ca/ipt/resource?r=cbg_canadian_specimens&v=1.4). The registered occurrence dataset can also be accessed directly from GBIF‘s web portal (https://doi.org/10.15468/mbwnw9), where it is available under a CC-BY 4.0 license (Data Citation 3). This release on GBIF extends exposure for the occurrence data and supports the Convention on Biological Diversity (CBD) by providing a dataset useful for its assessments and indicators [64], such as the Intergovernmental Science-Policy Platform on Biodiversity and Ecosystem Services (IPBES).

The DNA extracts derived from the 1,500,003 barcoded specimens are held in the DNA Archive at the CBG, either within −80°C freezers or dried in a trehalose-based preservative and held in −20°C freezers. The specimen data and DNA storage information were submitted to the Global Genome Biodiversity Network (GGBN) Data Portal [65] following the GGBN Data Standard [66]. The upload of CBG’s DNA extract data added 566 families and 4,287 genera to GGBN, increasing the number of families and genera by 22% and 30%, respectively (based on an API download of Animalia records from GGBN in May 2019). The data are accessible – and the DNA extracts can be requested on a cost-recovery basis through the GGBN portal (http://www.ggbn.org/ggbn_portal/search/result?institution=BIOUG%2C+Guelph) or University of Guelph’s IPT (https://ipt.uoguelph.ca/ipt/resource?r=public_data&v=1.8) (Data Citation 4).

## Technical Validation

### Inclusion in Data Release

Following taxonomic and sequence curation, specimens were required to pass one of two criteria before inclusion in the release dataset. First, specimens assigned to a BIN by the RESL algorithm (see Methods) were included. Second, if a specimen did not receive a BIN assignment, it was included in the dataset as long as its sequence was at least 300 bp long with <1% ambiguous base pairs, and led at least to a family-level assignment. No specimen whose sequence record was contaminated, had a stop codon, or was flagged by a member of the BOLD community (see Taxonomic Validation below) was included in the dataset.

### Sample Tracking

Using a custom-built collection information management system (CIMS), the specific location and storage medium for each specimen was captured at the time of its submission to the CBG’s collection archive. Unlike most natural history collections, specimens are arranged in order of processing to permit rapid submission of new specimens (up to 40,000 per week), to facilitate specimen retrieval (e.g., for photography), and to optimize the use of cabinet space. Because every specimen in the archive is databased, it is possible to query the CIMS (e.g., by a list of BINs, or a taxon for a particular geographical area) and quickly assemble all specimens required for an external loan or for examination by a visiting researcher.

### Taxonomic Validation

Multiple curatorial efforts were undertaken to validate taxonomic assignments. Taxonomic conflicts within BINs were investigated and resolved where possible. This review often led to a persistent flag in BOLD stating that the record is contaminated or misidentified (which works much like a wiki – see Pennisi 2008). The list of matches provided by the BOLD-ID Engine was checked for taxonomic discordances indicative of contaminated samples or misidentified specimens and corresponding data records were flagged. Neighbor-Joining (NJ) trees of similar taxa (at the order level) were constructed on BOLD to reveal unexpected placements of taxa; this also included evaluation of an image library paired to the tree to facilitate the recognition of specimens whose phenotype was incongruent with its taxonomic assignment (see Supplementary Files 3 and 4 for an example NJ tree and associated images for Canadian net-winged insects, Neuroptera).

All species-level identifications were validated against current nomenclature. The first validation pass included comparisons against national or regional checklists (e.g., [68] for true bugs; [69] for beetles; and [70] for moths and butterflies). Taxa that did not match with authoritative checklists were verified against online resources such as the Catalog of Life, WoRMs, ITIS, GBIF, or the World Spider Catalog. Remaining names were searched on a case-by-case basis in the taxonomic literature. Any synonyms or misspellings that were detected were corrected to the valid name.

### Sequence Validation

DNA sequences submitted to BOLD are first translated into amino acids and are then compared against a Hidden Markov Model of the COI protein. This pre-screening identifies gaps that provoke a frameshift or a stop codon, and other sequencing or editing errors. Sequences found to possess potential errors were manually re-edited or re-assembled from chromatogram trace files in CodonCode Aligner which often enabled the correction of errors made during the initial sequence editing. Sequences with confirmed gaps leading to frameshifts were excluded from the dataset. After initial submission to NCBI, staff at GenBank would report any residual errors detected with their validation tools allowing their correction before final submission.

## Conclusion

The DNA barcode reference library presented here, covering nearly 65,000 species of Canadian invertebrates, should have wide utility in supporting specimen identifications through barcoding and metabarcoding. Its primary use will undoubtedly derive from its capacity to assign unknown specimens and samples to a taxon. This step is key in producing accurate and reproducible data in metabarcoding studies [71,72]. The present DNA barcode reference library should also aid in quality control and validation for whole genome analysis by detecting misidentified samples and revealing cases of contamination (e.g. [73]). While the library will be most useful for work in Canada, a third of the species found in the Nearctic occurs in Canada, and about 5% of the Holarctic fauna, meaning the library will have utility across the Holarctic region. In fact, given its taxonomic breadth – 14 phyla, 43 classes, 163 orders, and 1123 families – it should be useful for studies worldwide, particularly for terrestrial invertebrates. It should also be valuable as a model for library construction in other countries and for other environments (e.g. soils, oceans), in Canada and elsewhere. In all applications, the accessibility of the library in various repositories (Data Citations 1-4), paired with the ongoing curation and refinement of taxonomic assignments by the biodiversity science community, further ensures its value will increase through time.

## Acknowledgements

The synthesis of this dataset was enabled by funding from the Canada Foundation for Innovation, from Genome Canada through Ontario Genomics, from NSERC, and from the Ontario Ministry of Research, Innovation and Science in support of the International Barcode of Life project. It was also enabled by philanthropic support from the Gordon and Betty Moore Foundation and from Ann McCain Evans and Chris Evans. The release of the data on GGBN was supported by a GGBN – Global Genome Initiative Award and we thank G. Droege, L. Loo, K. Barker, and J. Coddington for their support. Our work depended heavily on the analytical capabilities of the Barcode of Life Data Systems (BOLD, www.boldsystems.org). We also thank colleagues at the CBG for their support, including S. Adamowicz, S. Bateson, E. Berzitis, V. Breton, V. Campbell, A. Castillo, C. Christopoulos, J. Cossey, C. Gallant, J. Gleason, R. Gwiazdowski, M. Hajibabaei, R. Hanner, K. Hough, P. Janetta, A. Pawlowski, S. Pedersen, J. Robertson, D. Roes, K. Seidle, M. A. Smith, B. St. Jacques, A. Stoneham, J. Stahlhut, R. Tabone, J. Topan, S. Walker, C. Wei and T. Zemlak. For bioblitz-related assistance, we are grateful to D. Ireland, D. Metsger, A. Guidotti, J. Quinn and other members of Bioblitz Canada and Ontario Bioblitz. For our work in Canada’s national parks, we thank S. Woodley and J. Waithaka for their lead role in organizing permits and for the many Parks Canada staff who facilitated specimen collections, including M. Allen, D. Amirault-Langlais, J. Bastick, C. Belanger, C. Bergman, J.-F. Bisaillon, S. Boyle, J. Bridgland, S. Butland, L. Cabrera, R. Chapman, J. Chisholm, B. Chruszcz, D. Crossland, H. Dempsey, N. Denommee, T. Dobbie, C. Drake, J. Feltham, A. Forshner, K. Forster, S. Frey, L. Gardiner, P. Giroux, T. Golumbia, D. Guedo, N. Guujaaw, S. Hairsine, E. Hansen, C. Harpur, S. Hayes, J. Hoffman, D. Iles, S. Irwin, B. Johnston, V. Kafka, N. Kang, P. Langan, P. Lawn, M. Mahy, D. Masse, D. Mazerolle, C. McCarthy, I. McDonald, J. McIntosh, C. McKillop, V. Minelga, C. Ouimet, S. Parker, N. Perry, J. Piccin, A. Promaine, P. Roy, M. Savoie, D. Sigouin, P. Sinkins, R. Sissons, C. Smith, R. Smith, H. Stewart, G. Sundbo, D. Tate, R. Thompson, E. Tremblay, Y. Troutet, K. Tulk, J. Van Wieren, C. Vance, G. Walker, D. Whitaker, C. White, R. Wissink, C. Wong, and Y. Zharikov. Many other organizations improved coverage in the reference library by providing access to specimens – they included the Canadian National Collection of Insects, Arachnids and Nematodes, Smithsonian Institution’s National Museum of Natural History, the Canadian Museum of Nature, the University of Guelph Insect Collection, the Royal British Columbia Museum, the Royal Ontario Museum, the Pacific Forestry Centre, the Northern Forestry Centre, the Lyman Entomological Museum, the Churchill Northern Studies Centre, and *rare* Charitable Research Reserve. We also thank the many taxonomic specialists who identified specimens, including A. Borkent, B. Brown, M. Buck, C. Carr, T. Ekrem, J. Fernandez Triana, C. Guppy, K. Heller, J. Huber, L. Jacobus, J. Kjaerandsen, J. Klimaszewski, D. Lafontaine, J-F. Landry, G. Martin, A. Nicolai, D. Porco, H. Proctor, D. Quicke, J. Savage, B. C. Schmidt, M. Sharkey, A. Smith, E. Stur, A. Thomas, J. Webb, N. Woodley, and X. Zhou. We also thank K. Kerr and T. Mason for facilitating collections at Toronto Zoo and D. Iles for servicing the trap at Wapusk National Park.

This paper contributes to the University of Guelph’s Food from Thought research program supported by the Canada First Research Excellence Fund.

## Author Contributions

P.D.N.H., J.R.D., S.R., E.V.Z., A.V.B., and D.S. conceived and designed the experiments, including specimen processing and analytical pipelines. P.D.N.H. acquired the funding required to complete the work. K.P., J.E.S., M.R.Y., V.L.-B., C.N.S., G.B., K.K.S.L., J.T.A.M., R.M., M.P., A.E.R. and C.W. executed the field sampling and processed the specimens. A.A., S.L.D., N.V.I., L.L., N.M., S.N., N.N., and S.W.J.P performed the laboratory work. K.B., C.H., R.M., M.A.M., and C.S. provided informatics support. A.C.T, M.R.Y., J.E.S., and S.W.J.P. analyzed the data. J.R.D. led development of the manuscript, while P.D.N.H., A.V.B., S.R., D.S., and E.V.Z. provided substantive edits. All other authors read and approved the final manuscript.

## Data Citations

1. deWaard, J. R. et al. *Zenodo*. https://doi.org/10.5281/zenodo.3250923 (2019).
2. *NCBI BioProject* PRJNA472144 (2018).
3. BIOUG Centre for Biodiversity Genomics - Canadian Specimens. University of Guelph. https://doi.org/10.15468/mbwnw9 (2018).
4. BIOUG Centre for Biodiversity Genomics – DNA for Canadian Specimens. University of Guelph. https://doi.org/10.15468/mbwnw9 (2018).

## Supplementary Information

1. Duration of sampling from 2012-2014 as visualized by the number of Julian days of collecting for the ‘National Parks’ subset.
2. Breakdown of the methods used in the five major collection programs
3. Example BOLD Neighbor-Joining tree for Canadian net-winged insects (Neuroptera).
4. Example BOLD image library (associated with File 5) for Canadian net-winged insects (Neuroptera).
5. Taxonomic breakdown for the eight major collection methods used in the ‘National Parks’ subset. Taxonomic breakdown is summarized for A) specimens and B) BINs captured.
6. Summary data for the ‘National Parks’ subset.
7. Summary data for the ‘Other Localities’ subset.

## References

1. Cristescu, M. E. From barcoding single individuals to metabarcoding biological communities. Trends Ecol Evol 29, 566–571 (2014).

2. Deiner, K. et al. Environmental DNA metabarcoding: Transforming how we survey animal and plant communities. Mol Ecol 26, 5872–5895 (2017).

3. Braukmann, T. W. A. et al. Metabarcoding a diverse arthropod mock community. Mol Ecol Res 19, 711–727 (2019).

4. Taberlet, P., Coissac, E., Hajibabaei, M. & Rieseberg, L. H. Environmental DNA. Mol Ecol 21, 1789–1793 (2012).

5. Epp, L. S. et al. New environmental barcodes for analysing soil DNA: potential for studying past and present ecosystems. Mol Ecol 21, 1821–1833 (2012).

6. Hebert, P. D. N. et al. A Sequel to Sanger: Amplicon sequencing that scales. BMC Genomics 19, 14 (2018).

7. Benson, D. A., Karsch-Mizrachi, I., Lipman, D. J., Ostell, J. & Wheeler, D. L. GenBank. Nucl Acids Res 36, D25–D30 (2008).

8. Ratnasingham, S. & Hebert, P. D. N. BOLD: The Barcode of Life Data System (www.barcodinglife.org). Mol Ecol Notes 7, 355–364 (2007).

9. Machida, R. J., Leray, M., Ho, S.-L. & Knowlton, N. Metazoan mitochondrial gene sequence reference datasets for taxonomic assignment of environmental samples. Sci Data 4, 170027 (2017).

10. Heller, P., Casaletto, J., Ruiz, G. & Geller, J. B. A database of metazoan cytochrome c oxidase subunit I gene sequences derived from GenBank with CO-ARBitrator. Sci Data 5, 180156 (2018).

11. Ekrem, T., Willassen, E. & Stur, E. A comprehensive DNA sequence library is essential for identification with DNA barcodes. Mol Phylogenet Evol 43, 530–542 (2007).

12. Wilson, J. J., et al. When species matches are unavailable are DNA barcodes correctly assigned to higher taxa? An assessment using sphingid moths. BMC Ecol 11, 18 (2011).

13. Smit, J., Reijnen, B. T. & Stokvis, F. R. Half of the European fruit fly species barcoded (Diptera, Tephritidae); a feasibility test for molecular identification. ZooKeys 365, 279–305 (2013).

14. Coddington, J. A. et al. DNA barcode data accurately assign higher spider taxa. PeerJ 4, e2201 (2016).

15. Lou, M. & Golding, G. B. The effect of sampling from subdivided populations on species identification with DNA barcodes using a Bayesian statistical approach. Mol Phylogenet Evol 65, 765–773 (2012).

16. Meyer, C. P. & Paulay, G. DNA barcoding: Error rates based on comprehensive sampling. PLoS Biol 3, e422 (2005).

17. Kwong, S., Srivathsan, A. & Meier, R. An update on DNA barcoding: low species coverage and numerous unidentified sequences. Cladistics 28, 639–644 (2012).

18. Curry C. J., Gibson J. F., Shokralla, S., Hajibabaei, M. & Baird, D. J. Identifying North American freshwater invertebrates using DNA barcodes: are existing COI sequence libraries fit for purpose? Freshwater Sci 37, 178–189 (2018).

19. Zahiri, R. et al. 2017. Probing planetary biodiversity with DNA barcodes: The Noctuoidea of North America. PLoS ONE 12, e0178548 (2017).

20. Kvist, S. Barcoding in the dark? A critical view of the sufficiency of zoological DNA barcoding databases and a plea for broader integration of taxonomic knowledge. Mol Phylogenet Evol 69, 39–45 (2013).

21. Shokralla, S., et al. Next-generation DNA barcoding: Using next-generation sequencing to enhance and accelerate DNA barcode capture from single specimens. Mol Ecol Resour 14, 892–901 (2014).

22. Cruaud, P., Rasplus, J.-Y., Rodriguez, L.J. & Cruaud, A. High-throughput sequencing of multiple amplicons for barcoding and integrative taxonomy. Sci Rep 7, 41948 (2017).

23. Hebert, P. D. N., Cywinska, A., Ball, S. L. & deWaard, J. R. Biological identifications through DNA barcodes. Proc R Soc London Ser B 270, 313–321 (2003).

24. Prosser, S. W. J., deWaard, J. R., Miller, S. E., & Hebert, P. D. N. DNA barcodes from century-old type specimens using next-generation sequencing. Mol Ecol Resour 16, 487–497 (2016).

25. Ruedas, L. A., Salazar-Bravo, J., Dragoo, J. W. & Yates, T. L. The importance of being earnest: What, if anything, constitutes a “specimen examined?”. Mol Phylogenet Evol 17, 129–132 (2000).

26. Por, F. D. A “taxonomic affidavit”: Why it is needed? Integr Zool 2, 57–59 (2007).

27. Floyd, R., Lima, J., deWaard, J. R., Humble, L. R. & Hanner, R. H. Common goals: incorporating DNA barcoding into international protocols for identification of arthropod pests. Biol Invasions 12, 2947–2954 (2010).

28. Miller, S. E., Hausmann, A., Hallwachs, W. & Janzen, D. H. Advancing taxonomy and bioinventories with DNA barcodes. Philos Trans R Soc London Ser B 371, 20150339 (2016).

29. Bortolus, A. Error cascades in the biological sciences: The unwanted consequences of using bad taxonomy in ecology. Ambio 37, 114–118 (2008).

30. Groenenberg, D. S. J., Neubert, E. & Gittenberger, E. Reappraisal of the “Molecular phylogeny of Western Palaearctic Helicidae s.l. (Gastropoda: Stylommatophora)”: When poor science meets GenBank. Mol Phylogenet Evol 61, 914–923 (2011).

31. Monk, R. R. & Baker, R. J. e-Vouchers and the use of digital imagery in natural history collections. Museology 10, 1–8 (2001).

32. Hanner, R. H. & Gregory, T. R. Genomic diversity research and the role of biorepositories. Cell Preserv Technol 5, 93–103 (2007).

33. González, V. L. et al. Open access genomic resources for terrestrial arthropods. Curr Opin Insect Sci 25, 91–98 (2018).

34. Coissac, E., Hollingsworth, P. M., Lavergne, S. & Taberlet, P. 2016 From barcodes to genomes: extending the concept of DNA barcoding. Mol Ecol 25, 1423–1428 (2016).

35. Lewin, H. A. et al. Earth BioGenome Project: Sequencing life for the future of life. Proc Natl Acad Sci USA 115, 4325–4333 (2018).

36. Federal, Provincial and Territorial Governments of Canada. Canadian Biodiversity: Ecosystem Status and Trends 2010. Canadian Councils of Resource Ministers. Ottawa, ON, vi + 142 pp. (2010).

37. Malaise, R. A new insect-trap. Entomol Tidskr 58, 148–160 (1937).

38. Townes, H. Design for a Malaise trap. Proc. Entomol. Soc. Washington 64, 253–262 (1962).

39. Marston, N. Recent modifications in the design of Malaise traps with a summary of the insects represented in the collections. Journal of Kansas Entomological Society 38, 154–162 (1965).

40. Perez, K. H., Sones, J. E., deWaard, J. R. & Hebert, P. D. N. Mapping terrestrial biodiversity across the planet: a progress report on the Global Malaise Program. Genome, 60, 983 (2017).

41. Marshall, S. A., Anderson, R. S., Roughley, R. E., Behan-Pelletier, V. & Danks, H. V. Terrestrial arthropod biodiversity: planning a study and recommended sampling techniques. A brief prepared by the Biological Survey of Canada (Terrestrial Arthropods). Bull Entomol Soc Can 26, Supplement, 1–33 (1994).

42. Steinke, D., Breton, V., Berzitis, E. & Hebert, P. D. N. The School Malaise Trap Program: Coupling educational outreach with scientific discovery. PLoS Biol 15, e2001829 (2017).

43. Lundmark, C. BioBlitz: Getting into backyard biodiversity. BioScience 53, 329 (2003).

44. Zhou, X., Adamowicz, S. J., Jacobus, L. M., DeWalt, R. E. & Hebert, P. D. N. Towards a comprehensive barcode library for arctic life - Ephemeroptera, Plecoptera, and Trichoptera of Churchill, Manitoba, Canada. Front Zool 6, 30 (2009).

45. Young, M., Behan-Pelletier, V. & Hebert, P. D. N. Revealing the hyperdiverse mite fauna of subarctic Canada through DNA barcoding. PLoS ONE 7, e48755 (2012).

46. Woodcock, T. S. et al. The diversity and biogeography of the Coleoptera of Churchill: Insights from DNA barcoding. BMC Ecol 13, 40 (2013).

47. Blagoev, G. A., Nikolova, N. I., Sobel, C. N., Hebert, P. D. N. & Adamowicz, S.J. Spiders (Araneae) of Churchill, Manitoba: DNA barcodes and morphology reveal high species diversity and new Canadian records. BMC Ecology 13, 44 (2013).

48. Ratnasingham, S. & Hebert, P. D. N. BOLD’s role in barcode data management and analysis: A response. Mol Ecol Res 11, 941–942 (2011).

49. R Core Team. R: A Language and Environment for Statistical Computing. Vienna, Austria, R Foundation for Statistical Computing. https://www.R-project.org/. (2018).

50. Chamberlain, S. bold: Interface to Bold Systems API. R package version 0.8.6. https://CRAN.R-project.org/package=bold (2018).

51. Borisenko, A. V., Sones J. E. & Hebert P. D. N. The front-end logistics of DNA barcoding: challenges and prospects. Mol Ecol Res 9, 27–34 (2009).

52. deWaard, J. R. et al. Expedited assessment of terrestrial arthropod diversity by coupling Malaise traps with DNA barcoding. Genome 62, 85–95 (2019).

53. Porco, D., Rougerie, R., Deharveng, L. & Hebert, P. D. N. Coupling non-destructive DNA extraction and voucher retrieval for small soft-bodied Arthropods in a high-throughput context: The example of Collembola. Mol Ecol Resour 10, 942–945 (2010).

54. Ivanova, N. V., deWaard, J. R., & Hebert, P. D. N. An inexpensive, automation-friendly protocol for recovering high-quality DNA. Mol Ecol Notes 6, 998–1002 (2006).

55. Ratnasingham, S. & Hebert, P. D. N. A DNA-based registry for all animal species: The Barcode Index Number (BIN) system. PLoS One 8, e66213 (2013).

56. Ivanova, N. V. & Kuzmina, M. L. Protocols for dry DNA storage and shipment at room temperature. Mol Ecol Res 13, 890–898 (2013).

57. Ding, W. & Taylor, G. Automatic moth detection from trap images for pest management. Comput Electron Agr 123, 17–28 (2016).

58. Mosquin, T., Whiting, P. G. & McAllister, D.E. Canada’s biodiversity: the variety of life, its status, economic benefits, conservation costs and unmet needs. Canadian Museum of Nature, Ottawa, ON. 293 pp. (1995).

59. Canadian Endangered Species Conservation Council. Wild Species 2015: The General Status of Species in Canada, National General Status Working Group, www.wildspecies.ca, (2016).

60. Langor, D. W. The diversity of terrestrial arthropods in Canada. In: Langor D. W. & Sheffield, C. S. (Eds) The Biota of Canada – A Biodiversity Assessment. Part 1: The Terrestrial Arthropods. ZooKeys 819: 9–40 (2019).

61. Song, H., Buhay, J. E., Whiting, M. F. & Crandall, K. A. Many species in one: DNA barcoding overestimates the number of species when nuclear mitochondrial pseudogenes are coamplified. Proc Natl Acad Sci USA 105, 13486–13491 (2008).

62. Wieczorek, J. et al. Darwin Core: An evolving community-developed biodiversity data standard. PLoS ONE 7, e29715 (2012).

63. Robertson T. et al. The GBIF Integrated Publishing Toolkit: Facilitating the efficient publishing of biodiversity data on the internet. PLoS ONE 9, e102623 (2014).

64. Vernooy, R. et al. Barcoding life to conserve biological diversity: Beyond the taxonomic imperative. PLoS Biol 8, e1000417 (2010).

65. Droege, G. et al. The Global Genome Biodiversity Network (GGBN) data portal. Nucl Acids Res 42, D607–D612 (2014).

66. Droege, G. et al. The Global Genome Biodiversity Network (GGBN) data standard specification. Database 2016, baw125 (2016).

67. Pennisi, E. DNA data. Proposal to ‘Wikify’ GenBank meets stiff resistance. Science 319, 1598–1599 (2008).

68. Maw, H. E. L., Foottit, R. G., Hamilton, K. G. A. & Scudder, G. G. E. Checklist of the Hemiptera of Canada and Alaska, Ottawa: NRC Research Press (2000).

69. Bousquet, Y., Bouchard, P., Davies, A. E. & Sikes, D.S. Checklist of beetles (Coleoptera) of Canada and Alaska. Second edition. ZooKeys 360, 1–402 (2013).

70. Pohl, G. R. et al. Annotated checklist of the moths and butterflies (Lepidoptera) of Canada and Alaska. Series Faunistica 118, 1–580 (2018).

71. Alsos, I. G. et al. Plant DNA metabarcoding of lake sediments: How does it represent the contemporary vegetation. PLoS ONE 13, e0195403 (2018).

72. Zinger, L. et al. DNA metabarcoding—Need for robust experimental designs to draw sound ecological conclusions. Mol Ecol 28, 1857–1862 (2019).

73. Espeland, M. et al. A comprehensive and dated phylogenomic analysis of butterflies. Curr Biol 28, 770–778 (2018).

