## Supplementary figures and images for "A reference library for the identification of Canadian invertebrates: 1.5 million DNA barcodes, voucher specimens, and genomic samples"

### Supplementary File 1

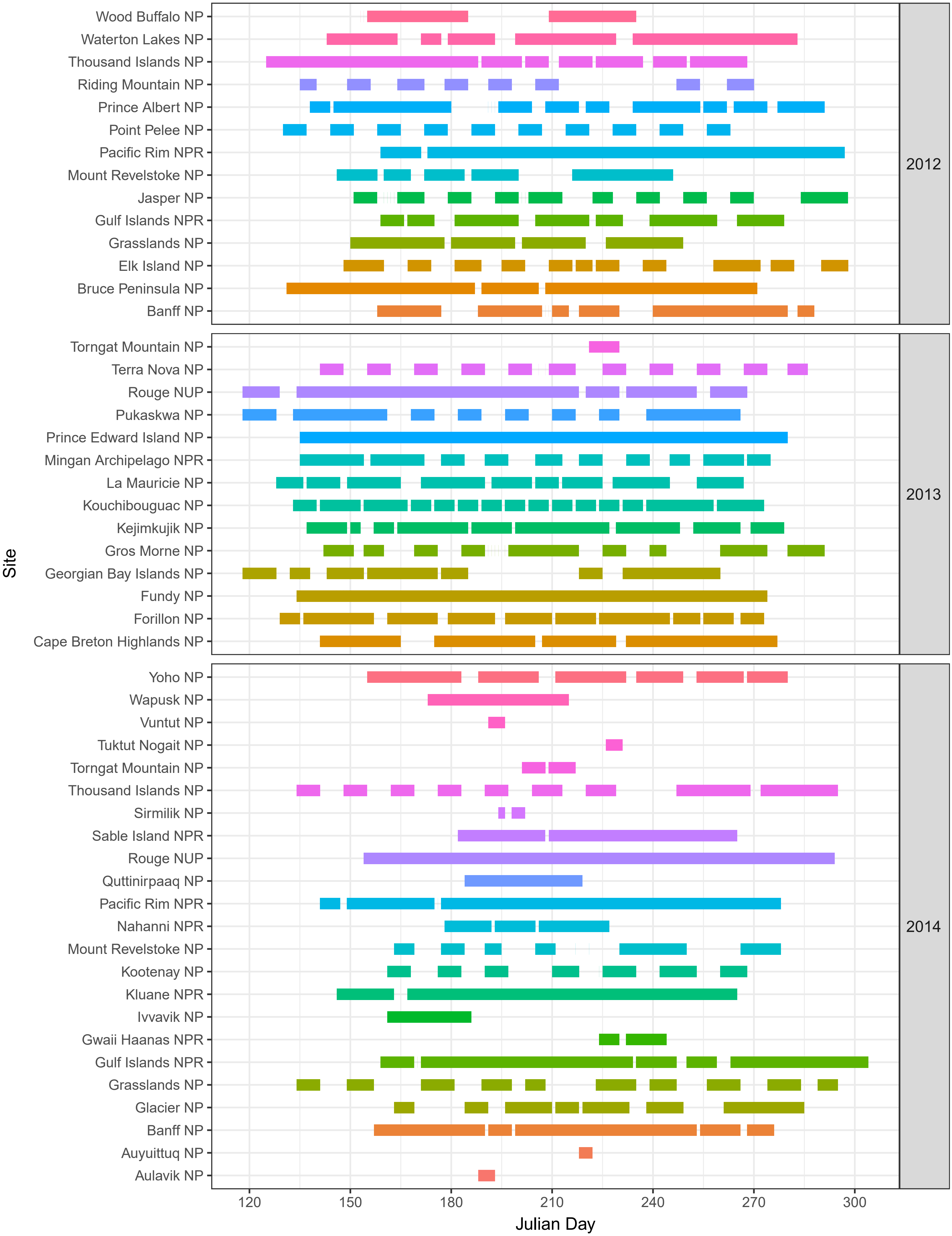

### Supplementary File 3

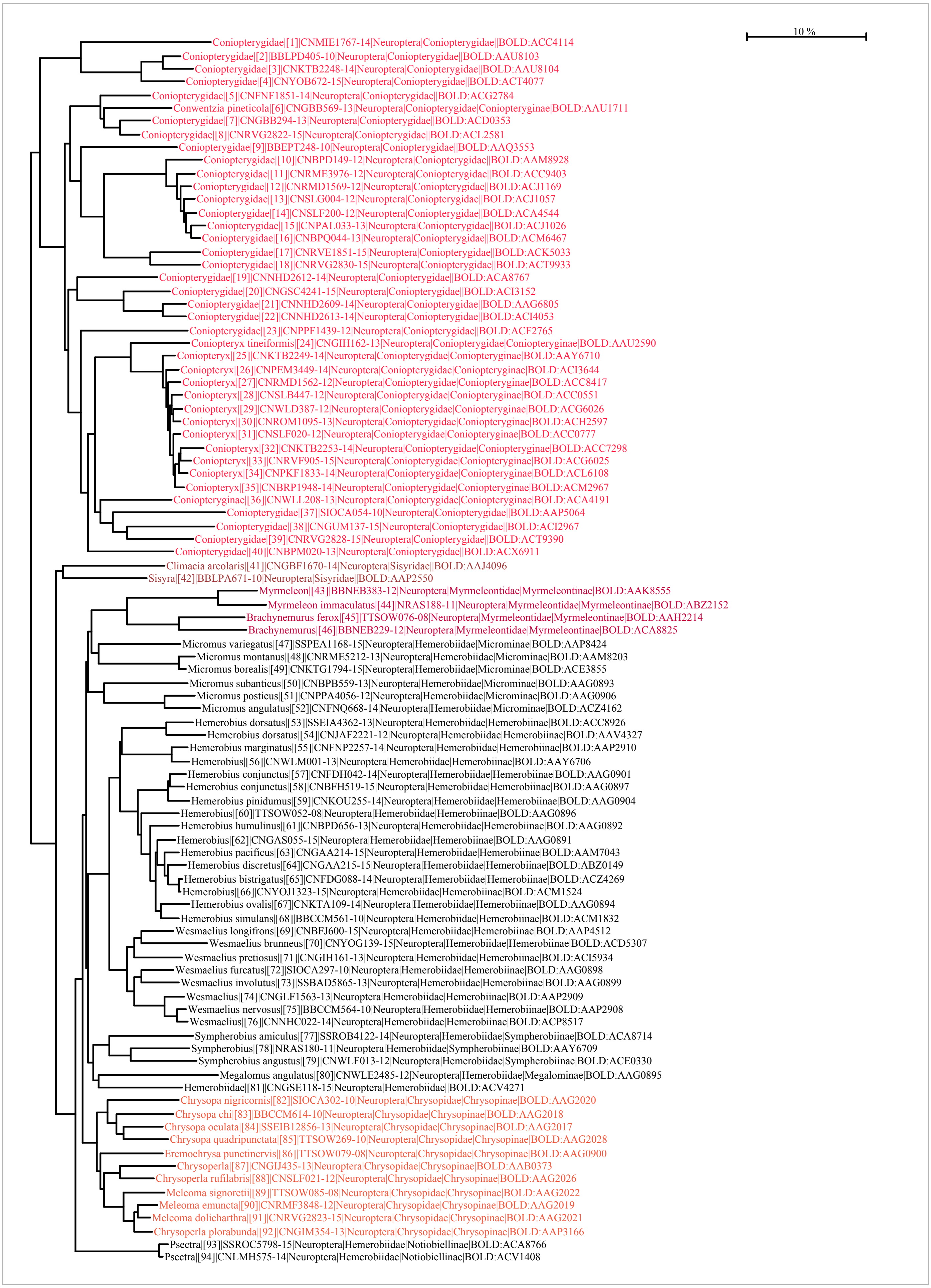

### Supplementary File 5

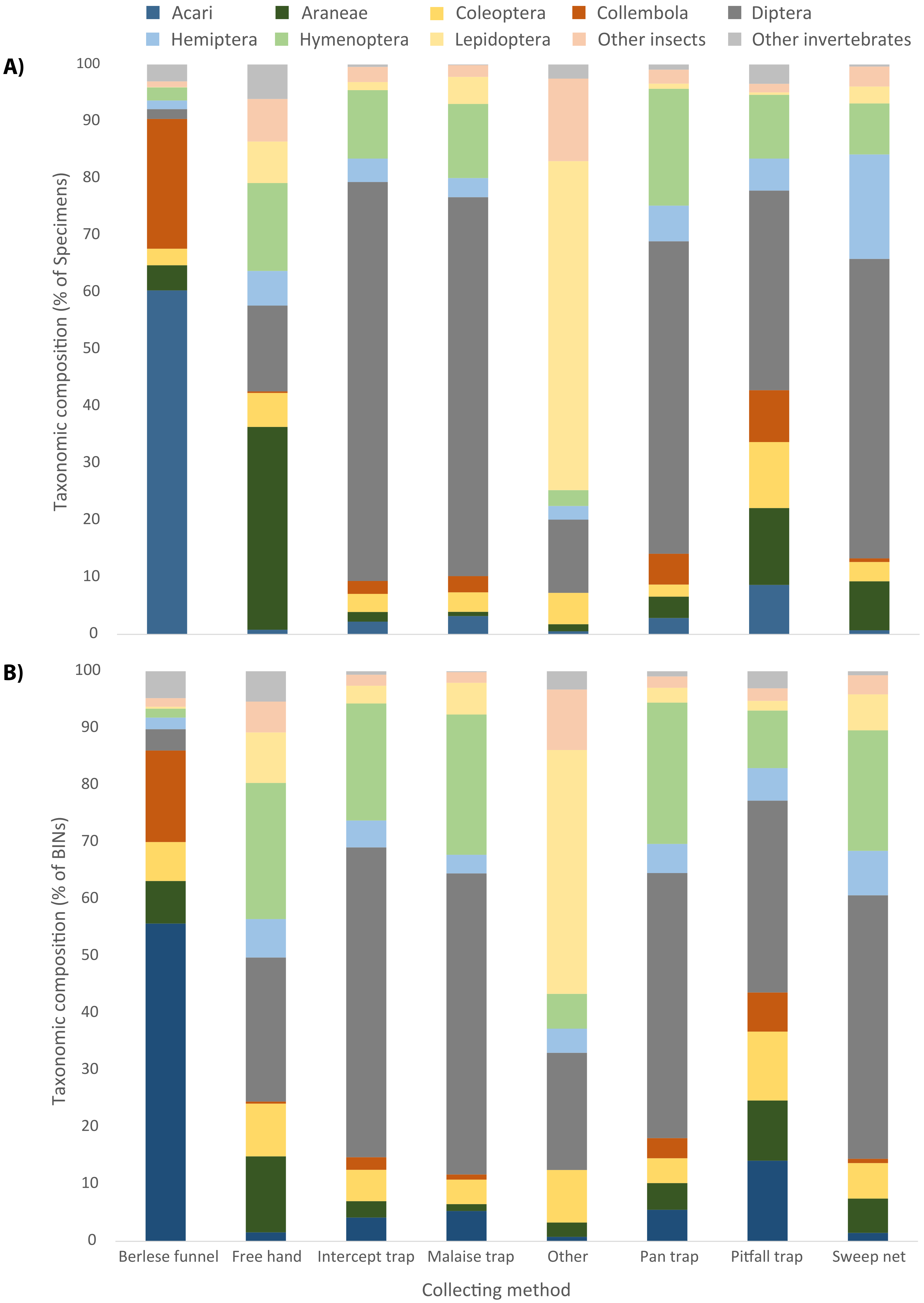
