## Supplementary File 2 for "A reference library for the identification of Canadian invertebrates: 1.5 million DNA barcodes, voucher specimens, and genomic samples"

| | A. National Parks Subset | | | B. Other Localities Subset | | | |
|---|---|---|---|---|---|---|---|
| | **Global Malaise Program - Canada** | **Standardized Sampling Program** | **Other Collections** | **Global Malaise Program - Canada** | **School Malaise Trap Program** | **All taxa biodiversity inventories and bioblitzes** | **Other Collections** |
| Malaise Trap | x | x | x | x | x | x | x |
| Intercept Trap | | x | x | | | x | x |
| Pan Trap | | x | x | | | x | x |
| UV Light Trap | | | x | | | x | x |
| UV Light Sheet | | | x | | | x | x |
| Pitfall trap | | x | x | | | x | x |
| Sweep Net | | x | x | | | x | x |
| Free Hand | | | x | | | x | x |
| Berlese Funnel | | | x | | | x | x |
| UV Bucket Trap | | | x | | | x | |
| Dip net | | | x | | | x | x |
| Bottle Trap | | | x | | | x | x |
| Mustard Extraction | | | x | | | | x |
| Sieve | | | x | | | | x |
| Plankton Net | | | x | | | x | x |
