## Supplementary File 4 for "A reference library for the identification of Canadian invertebrates: 1.5 million DNA barcodes, voucher specimens, and genomic samples"

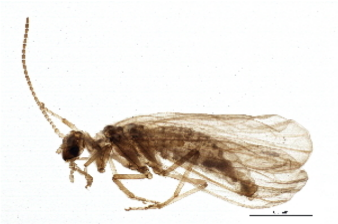

**BIOUG03259-H02 [Lateral]**
Coniopterygidae
BIN URI: BOLD:ACC4114

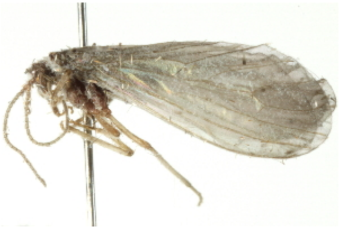

**BIOUG00762-B09 [Lateral]**
Coniopterygidae
BIN URI: BOLD:AAU8103

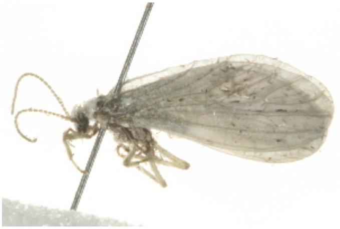

**BIOUG00762-D05 [Lateral]**
Coniopterygidae
BIN URI: BOLD:AAU8104

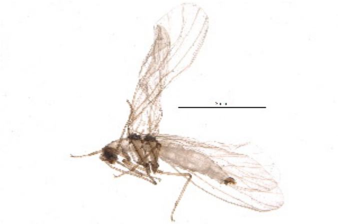

**BIOUG19021-H05 [Lateral]**
Coniopterygidae
BIN URI: BOLD:ACT4077

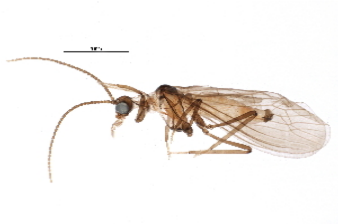

**BIOUG10477-A11 [Lateral]**
Coniopterygidae
BIN URI: BOLD:ACG2784

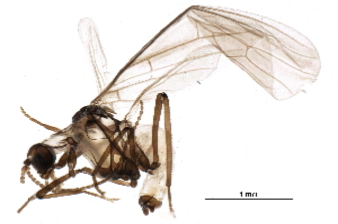

**BIOUG21944-D08 [Dorsal]**
Conwentzia pineticola
BIN URI: BOLD:AAU1711

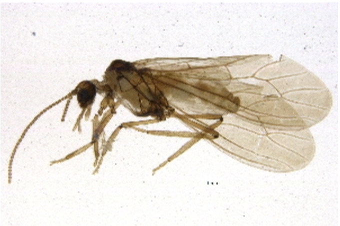

**BIOUG03911-E07 [Lateral]**
Coniopterygidae
BIN URI: BOLD:ACD0353

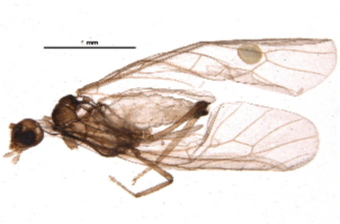

**BIOUG09967-G07 [Lateral]**
Coniopterygidae
BIN URI: BOLD:ACL2581

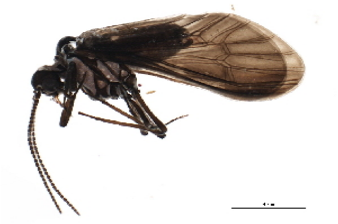

**10BBEPT-0247 [Lateral]**
Coniopterygidae
BIN URI: BOLD:AAQ3553

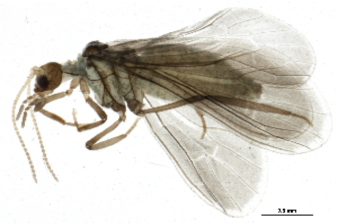

**BIOUG04493-G10 [Lateral]**
Coniopterygidae
BIN URI: BOLD:AAM8928

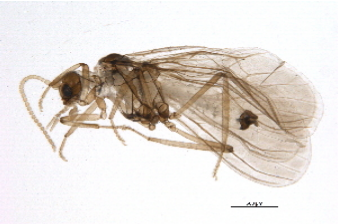

**BIOUG03727-A01 [Lateral]**
Coniopterygidae
BIN URI: BOLD:ACC9403

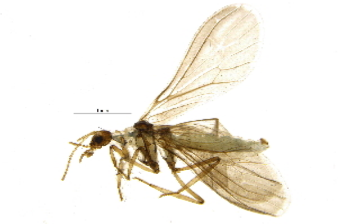

**BIOUG03022-F07 [Lateral]**
Coniopterygidae
BIN URI: BOLD:ACJ1169

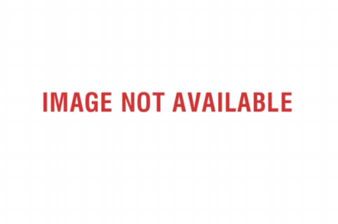


**BIOUG03080-H09**
Coniopterygidae
BIN URI: BOLD:ACJ1057

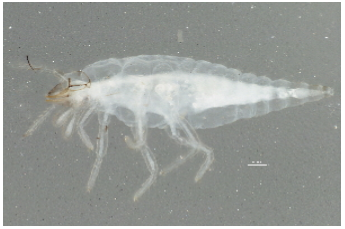

**BIOUG03716-H11 [Lateral]**
Coniopterygidae
BIN URI: BOLD:ACA4544

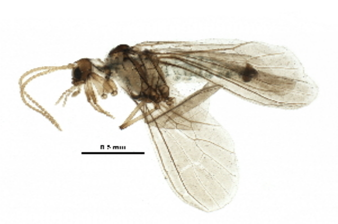

**BIOUG21153-H04 [Lateral]**
Coniopterygidae
BIN URI: BOLD:ACJ1026

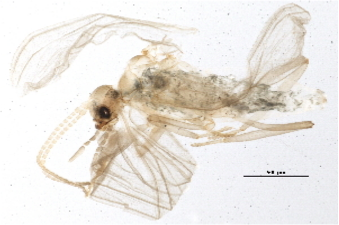

**BIOUG05981-F01 [Lateral]**
Coniopterygidae
BIN URI: BOLD:ACM6467

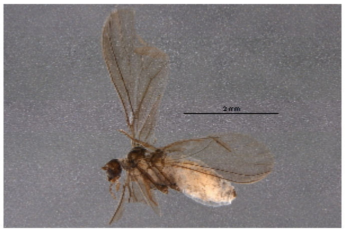
BIOUG07110-C06 [Lateral]
Coniopterygidae
BIN URI: BOLD:ACK5033

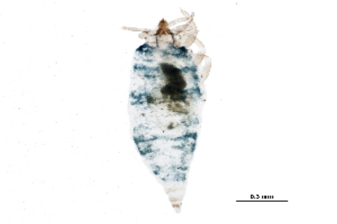
BIOUG20369-F05 [Dorsal]
Coniopterygidae
BIN URI: BOLD:ACT9933

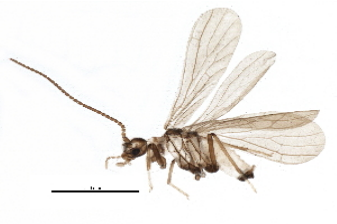
BIOUG17693-G11 [Lateral]
Coniopterygidae
BIN URI: BOLD:ACA8767

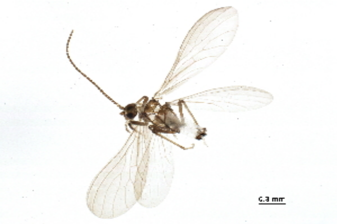
BIOUG07171-B08 [Lateral]
Coniopterygidae [skewed]
BIN URI: BOLD:ACI3152

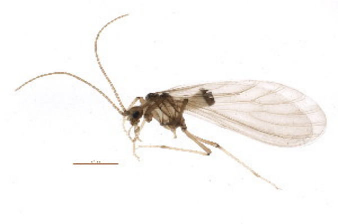
BIOUG03280-F10 [Lateral]
Coniopterygidae
BIN URI: BOLD:AAG6805

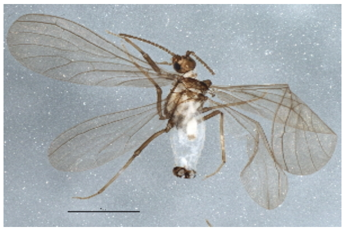
BIOUG06018-H10 [Dorsal]
Coniopterygidae
BIN URI: BOLD:ACI4053

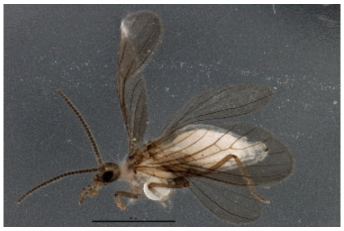
BIOUG03566-A09 [Lateral]
Coniopterygidae
BIN URI: BOLD:ACF2765

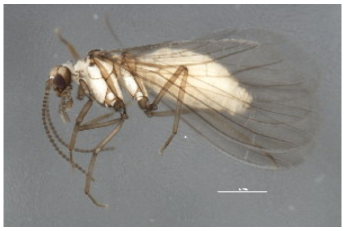
BIOUG00864-E03 [Lateral]
Coniopteryx tineiformis
BIN URI: BOLD:AAU2590

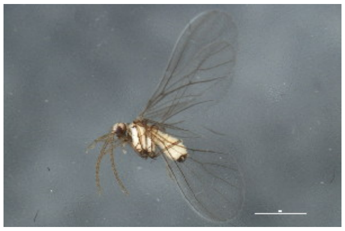
BIOUG00864-E04 [Lateral]
Coniopteryx
BIN URI: BOLD:AAY6710

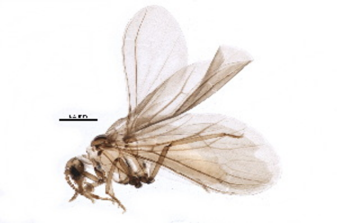
BIOUG11087-G10 [Lateral]
Coniopteryx
BIN URI: BOLD:ACI3644

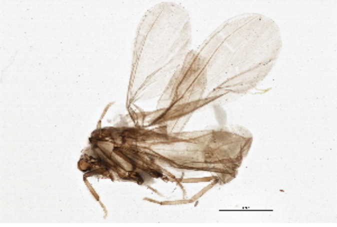
BIOUG03022-E12 [Lateral]
Coniopteryx
BIN URI: BOLD:ACC8417

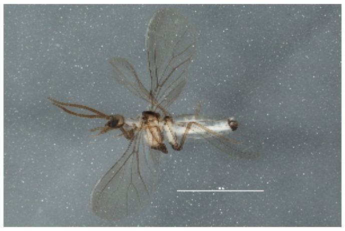
BIOUG02751-B10 [Lateral]
Coniopteryx
BIN URI: BOLD:ACC0551

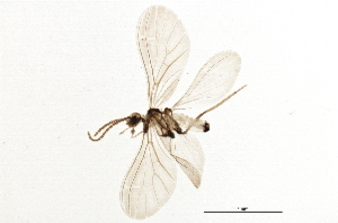
BIOUG03259-G04 [Lateral]
Coniopteryx
BIN URI: BOLD:ACG6026

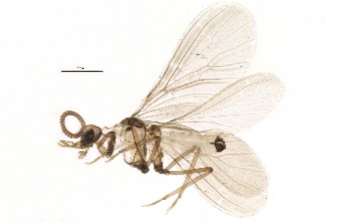
BIOUG06869-F09 [Lateral]
Coniopteryx
BIN URI: BOLD:ACH2597

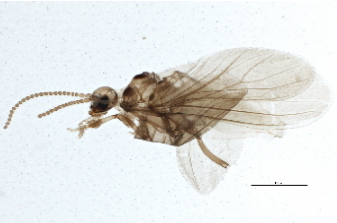
BIOUG04967-C05 [Lateral]
Coniopteryx
BIN URI: BOLD:ACC0777

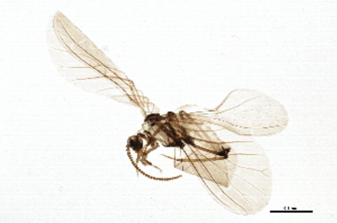
BIOUG03259-G06 [Lateral]
Coniopteryx
BIN URI: BOLD:ACC7298

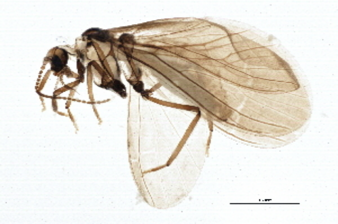
BIOUG03259-G07 [Lateral]
Coniopteryx
BIN URI: BOLD:ACG6025

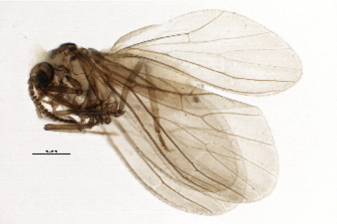
BIOUG10296-F02 [Lateral]
Coniopteryx
BIN URI: BOLD:ACL6108

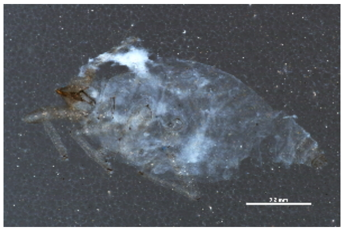
BIOUG11215-B08 [Lateral]
Coniopteryx
BIN URI: BOLD:ACM2967

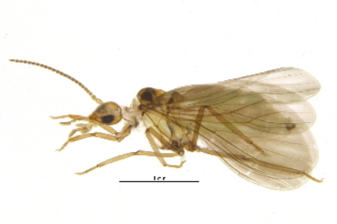
BIOUG03566-B01 [Lateral]
Coniopterygidae
BIN URI: BOLD:ACA4191

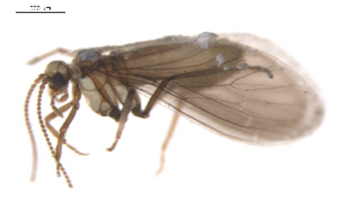
10BBSIO-0054 [Lateral]
Coniopterygidae
BIN URI: BOLD:AAP5064

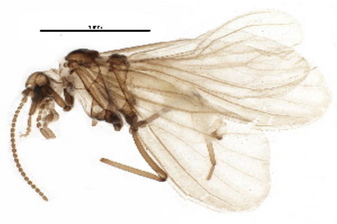
BIOUG06945-F06 [Lateral]
Coniopterygidae
BIN URI: BOLD:ACI2967

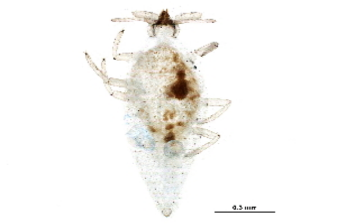
BIOUG20369-F03 [Dorsal]
Coniopterygidae
BIN URI: BOLD:ACT9390

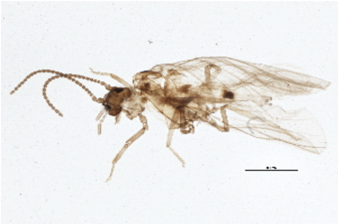
BIOUG05828-E10 [Lateral]
Coniopterygidae
BIN URI: BOLD:ACX6911

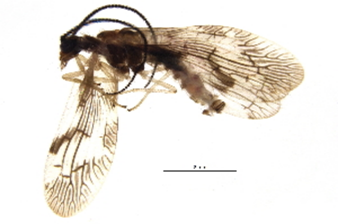

BIOUG10061-H03 [Lateral]
Climacia areolaris
BIN URI: BOLD:AAJ4096

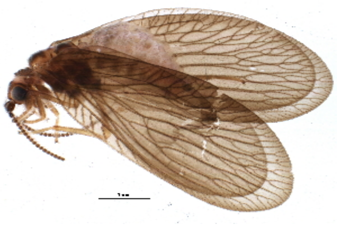

BIOUG08286-F11 [Lateral]
Sisyra
BIN URI: BOLD:AAP2550

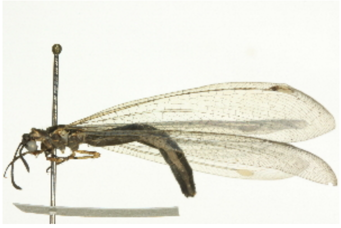

BIOUG02806-F02 [Lateral]
Myrmeleon
BIN URI: BOLD:AAK8555

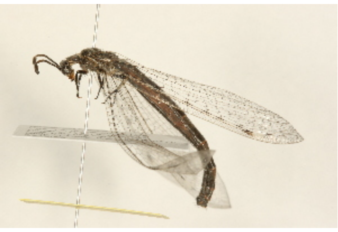

09BBNEU-0055 [Lateral]
Myrmeleon immaculatus
BIN URI: BOLD:ABZ2152

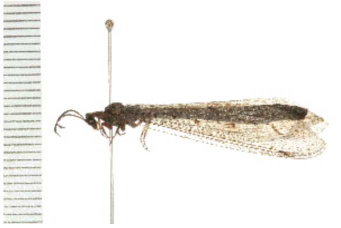

08BBNEU-076 [Lateral]
Brachynemurus ferox
BIN URI: BOLD:AAH2214

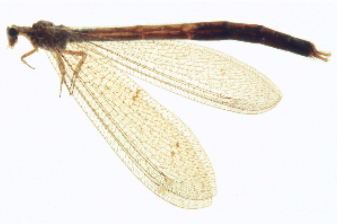

BIOUG02805-A01 [Lateral]
Brachynemurus
BIN URI: BOLD:ACA8825

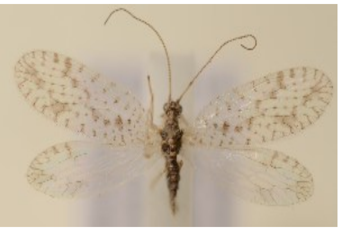

TR00040 [Dorsal] [skewed]
Micromus variegatus
BIN URI: BOLD:AAP8424

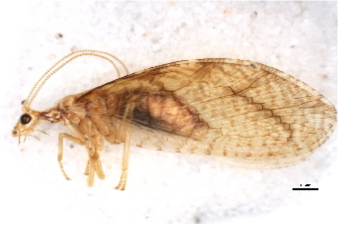

09BBNEU-0142 [Lateral]
Micromus montanus
BIN URI: BOLD:AAM8203

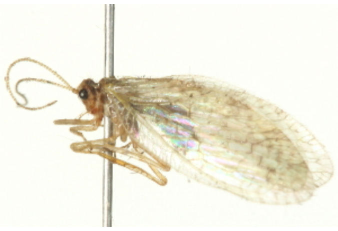

BIOUG00762-F11 [Lateral]
Micromus borealis
BIN URI: BOLD:ACE3855

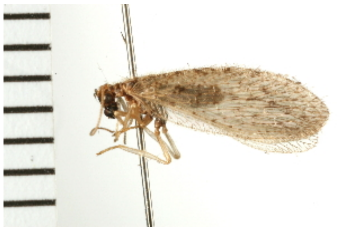

08BBNEU-013 [Lateral]
Micromus subanticus
BIN URI: BOLD:AAG0893

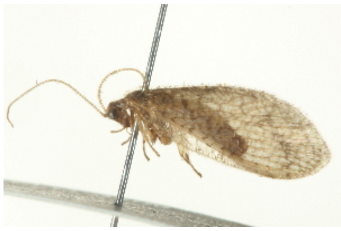

PHFLO-0106 [Lateral]
Micromus posticus
BIN URI: BOLD:AAG0906

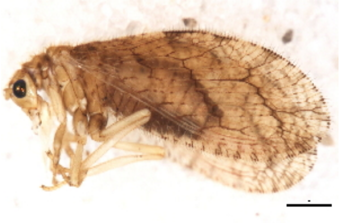

09BBNEU-0148 [Lateral]
Micromus angulatus
BIN URI: BOLD:ACZ4162

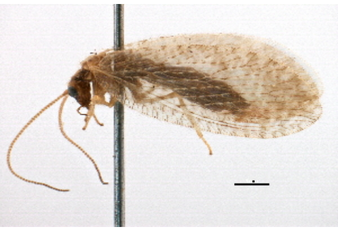

BIOUG10951-B11 [Lateral]
Hemerobius dorsatus
BIN URI: BOLD:ACC8926

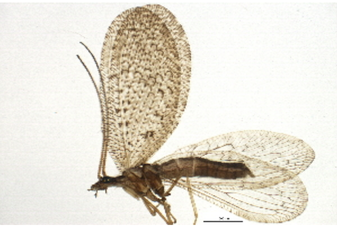

BIOUG03505-C06 [Lateral]
Hemerobius dorsatus
BIN URI: BOLD:AAV4327

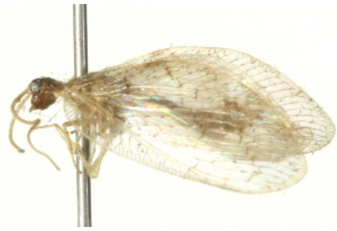

BIOUG00762-F03 [Lateral]
Hemerobius
BIN URI: BOLD:AAP2910

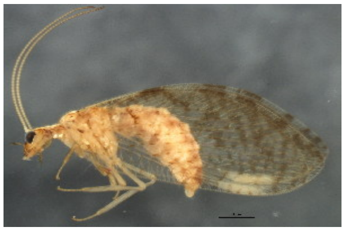

BIOUG00864-A06 [Lateral]
Hemerobius
BIN URI: BOLD:AAY6706

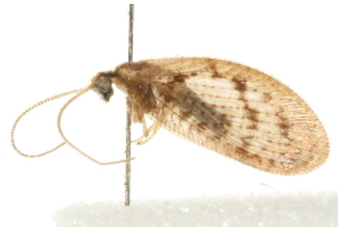

08BBNEU-088 [Lateral]
Hemerobius ovalis
BIN URI: BOLD:AAG0901

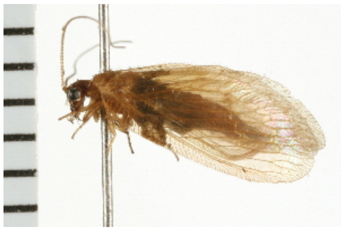

08BBNEU-063 [Lateral]
Hemerobius conjunctus
BIN URI: BOLD:AAG0897

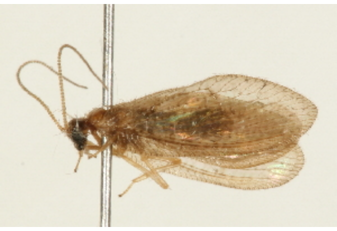

09BBNEU-0095 [Lateral]
Hemerobius pinidumus
BIN URI: BOLD:AAG0904

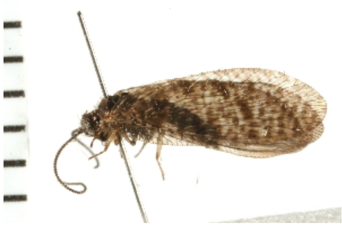

08BBNEU-052 [Lateral]
Hemerobius
BIN URI: BOLD:AAG0896

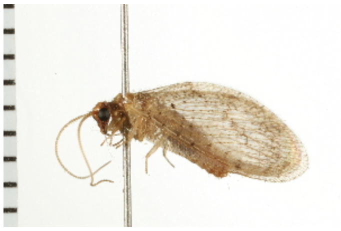

08BBNEU-005 [Lateral]
Hemerobius humulinus
BIN URI: BOLD:AAG0892

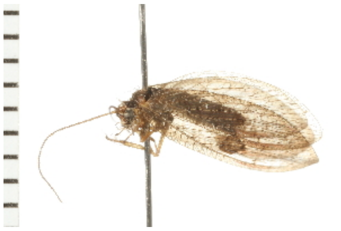

08BBNEU-018 [Lateral]
Hemerobius
BIN URI: BOLD:AAG0891

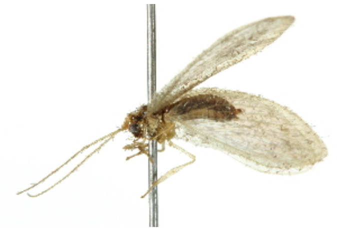

10BBNEU-0006 [Lateral]
Hemerobius
BIN URI: BOLD:AAM7043

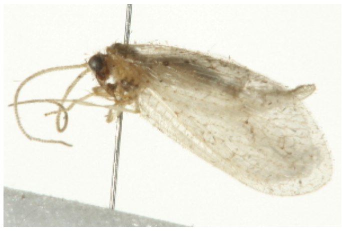

BIOUG00762-D07 [Lateral]
Hemerobius discretus
BIN URI: BOLD:ABZ0149

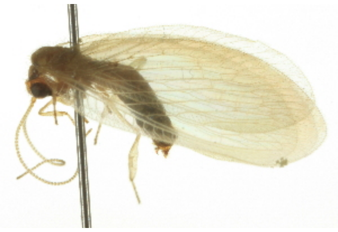

**BIOUG02795-A09 [Lateral]**
Hemerobius
BIN URI: BOLD:ACZ4269

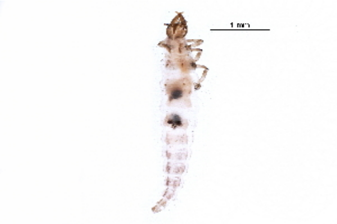

**BIOUG19485-B06 [Larva]**
Hemerobius
BIN URI: BOLD:ACM1524

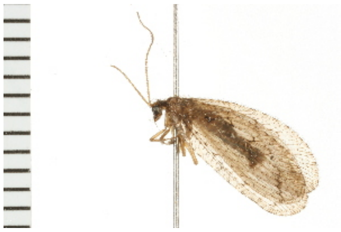

**08BBNEU-021 [Lateral]**
Hemerobius ovalis
BIN URI: BOLD:AAG0894

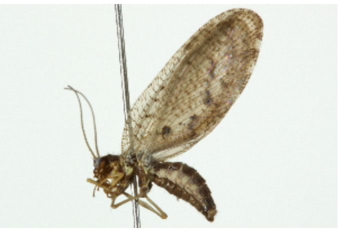

**09BBNEU-0169 [Lateral]**
Hemerobius simulans
BIN URI: BOLD:ACM1832

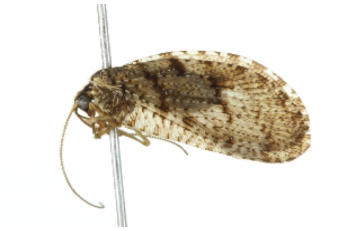

**09BBNEU-0173 [Lateral]**
Wesmaelius longifrons
BIN URI: BOLD:AAP4512

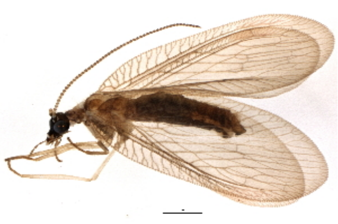

**BIOUG05803-G04 [Lateral]**
Wesmaelius brunneus
BIN URI: BOLD:ACD5307

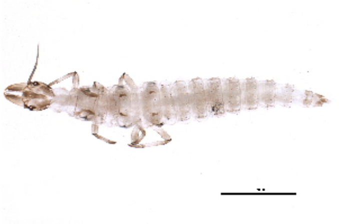

**BIOUG16124-G03 [Larva]**
Wesmaelius pretiosus
BIN URI: BOLD:ACI5934

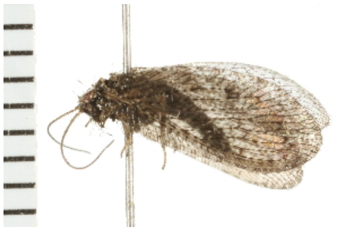

**08BBNEU-066 [Lateral]**
Wesmaelius furcatus
BIN URI: BOLD:AAG0898

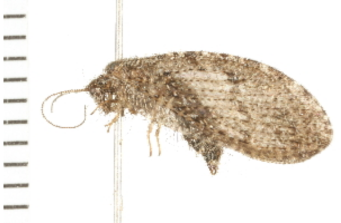

**08BBNEU-071 [Lateral]**
Wesmaelius involutus
BIN URI: BOLD:AAG0899

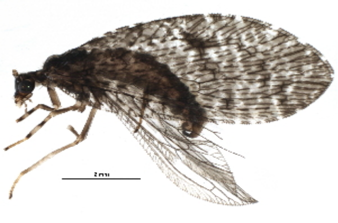

**BIOUG20171-E09 [Lateral]**
Wesmaelius
BIN URI: BOLD:AAP2909

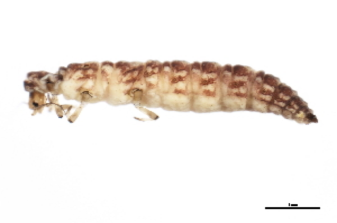

**BIOUG00676-A12 [Lateral]**
Wesmaelius nervosus
BIN URI: BOLD:AAP2908

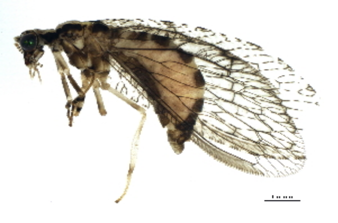

**BIOUG16073-F03 [Lateral]**
Wesmaelius
BIN URI: BOLD:ACP8517

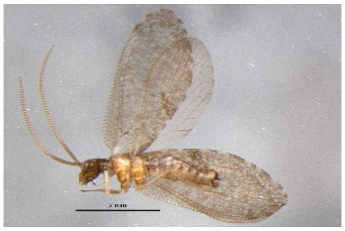

**BIOUG02863-D02 [Lateral]**
Sympherobius amiculus
BIN URI: BOLD:ACA8714

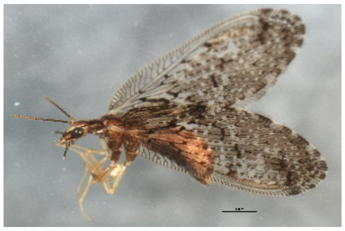

**BIOUG00864-D03 [Lateral]**
Sympherobius
BIN URI: BOLD:AAY6709

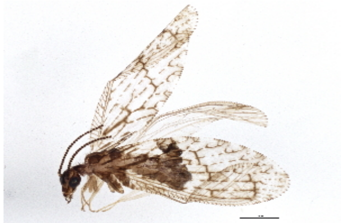

**BIOUG03522-C11 [Lateral]**
Sympherobius angustus
BIN URI: BOLD:ACE0330

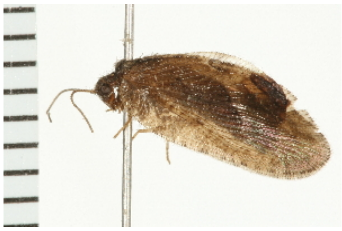

**08BBNEU-050 [Lateral]**
Megalomus angulatus
BIN URI: BOLD:AAG0895

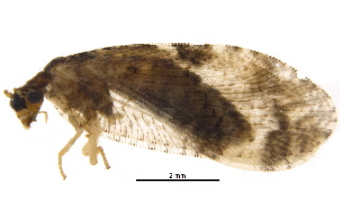

**BIOUG21523-F03 [Lateral]**
Hemerobiidae
BIN URI: BOLD:ACV4271

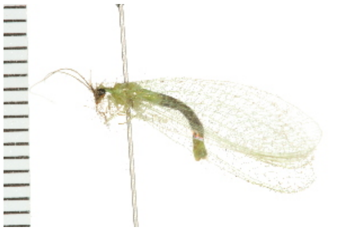

**08BBNEU-057 [Lateral]**
Chrysopa nigricornis
BIN URI: BOLD:AAG2020

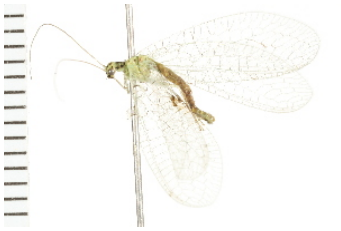

**08BBNEU-053 [Lateral]**
Chrysopa chi
BIN URI: BOLD:AAG2018

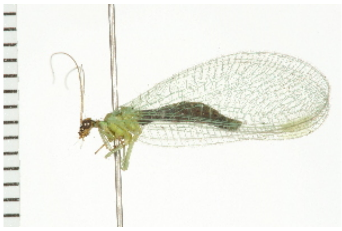

**08BBNEU-033 [Lateral]**
Chrysopa oculata
BIN URI: BOLD:AAG2017

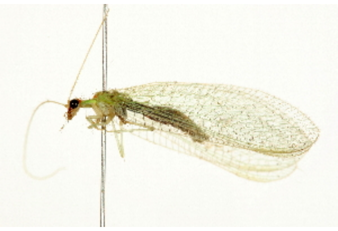

**09BBNEU-0003 [Lateral]**
Chrysopa quadripunctata
BIN URI: BOLD:AAG2028

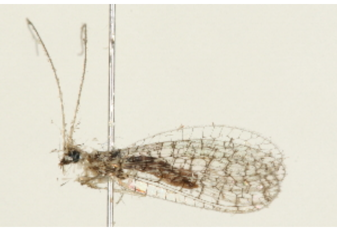

**09BBNEU-0097 [Lateral]**
Eremochrysa punctinervis
BIN URI: BOLD:AAG0900

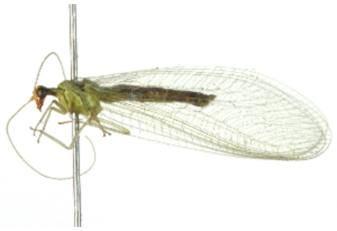

**10BBNEU-0012 [Lateral]**
Chrysoperla
BIN URI: BOLD:AAB0373

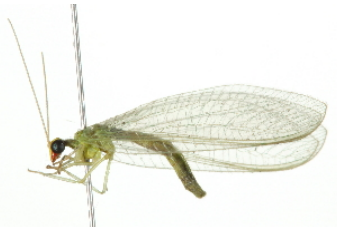

**10BBNEU-0033 [Lateral]**
Chrysoperla rufilabris
BIN URI: BOLD:AAG2026

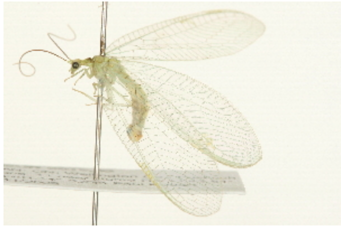

**NEUR 0019.02 [Lateral]**
Meleoma signoretii
BIN URI: BOLD:AAG2022

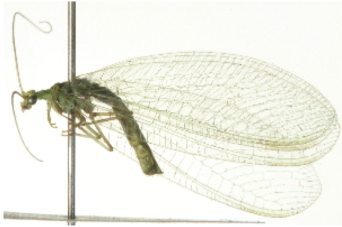

**BIOUG00762-B01 [Lateral]**
Meleoma emuncta
BIN URI: BOLD:AAG2019

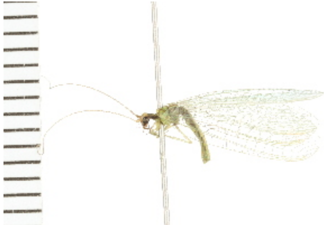

**08BBNEU-073 [Lateral]**
Meleoma dolicharthra
BIN URI: BOLD:AAG2021

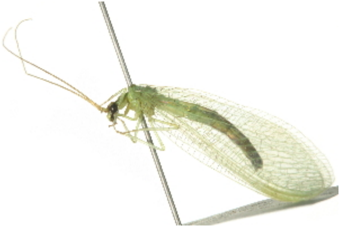

**BIOUG00762-F12 [Lateral]**
Chrysoperla plorabunda
BIN URI: BOLD:AAP3166

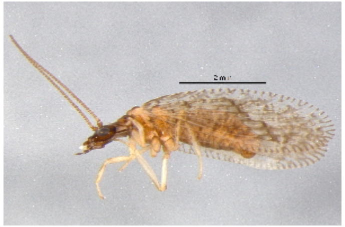

**BIOUG02863-D11 [Lateral]**
Psectra
BIN URI: BOLD:ACA8766

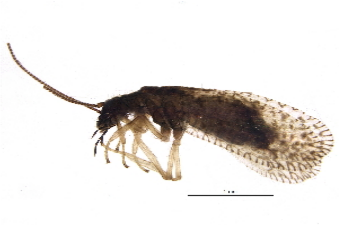

**BIOUG11001-E06 [Lateral]**
Psectra
BIN URI: BOLD:ACV1408
